# Mammalian deubiquitinating enzyme inhibitors display *in vitro* and *in vivo* activity against malaria parasites and potentiate artemisinin action

**DOI:** 10.1101/2020.08.13.249425

**Authors:** Nelson V. Simwela, Katie R. Hughes, Michael T. Rennie, Michael P. Barrett, Andrew P. Waters

## Abstract

Current malaria control efforts rely significantly on artemisinin combinational therapies which have played massive roles in alleviating the global burden of the disease. Emergence of resistance to artemisinins is therefore, not just alarming but requires immediate intervention points such as development of new antimalarial drugs or improvement of the current drugs through adjuvant or combination therapies. Artemisinin resistance is primarily conferred by Kelch13 propeller mutations which are phenotypically characterised by generalised growth quiescence, altered haemoglobin trafficking and downstream enhanced activity of the parasite stress pathways through the ubiquitin proteasome system (UPS). Previous work on artemisinin resistance selection in a rodent model of malaria, which we and others have recently validated using reverse genetics, has also shown that mutations in deubiquitinating enzymes, DUBs (upstream UPS component) modulates susceptibility of malaria parasites to both artemisinin and chloroquine. The UPS or upstream protein trafficking pathways have, therefore, been proposed to be not just potential drug targets, but also possible intervention points to overcome artemisinin resistance. Here we report the activity of small molecule inhibitors targeting mammalian DUBs in malaria parasites. We show that generic DUB inhibitors can block intraerythrocytic development of malaria parasites *in vitro* and possess antiparasitic activity *in vivo* and can be used in combination with additive effect. We also show that inhibition of these upstream components of the UPS can potentiate the activity of artemisinin *in vitro* as well as *in vivo* to the extent that ART resistance can be overcome. Combinations of DUB inhibitors anticipated to target different DUB activities and downstream 20s proteasome inhibitors are even more effective at improving the potency of artemisinins than either inhibitors alone providing proof that targeting multiple UPS activities simultaneously could be an attractive approach to overcoming artemisinin resistance. These data further validate the parasite UPS as a target to both enhance artemisinin action and potentially overcome resistance. Lastly, we confirm that DUB inhibitors can be developed into *in vivo* antimalarial drugs with promise for activity against all of human malaria and could thus further exploit their current pursuit as anticancer agents in rapid drug repurposing programs.

**Graphical abstract:** 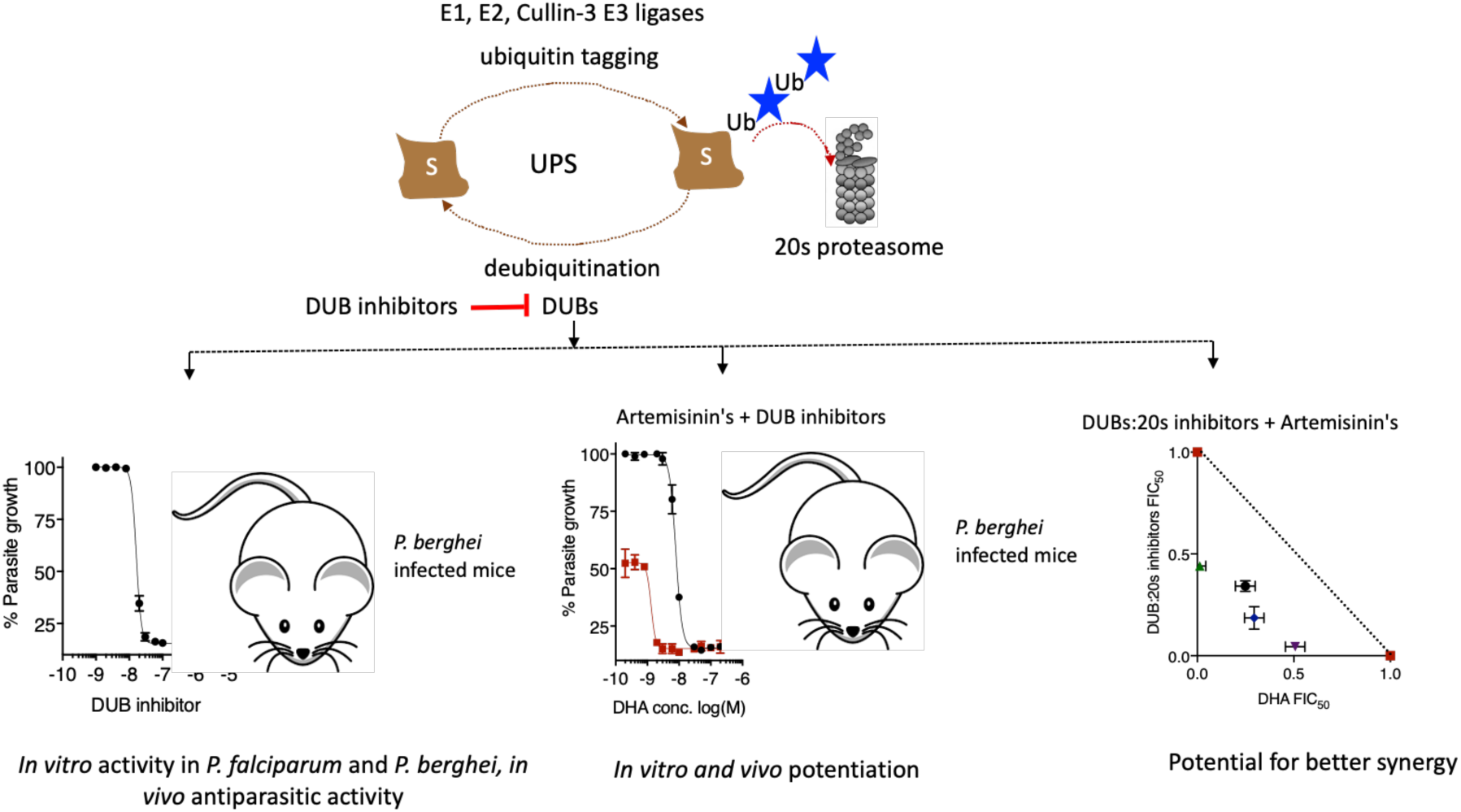

## Introduction

Malaria remains the most important parasitic disease in tropical and sub-tropical regions of the world with high rates of morbidity and mortality. Despite significant gains in malaria control over the past decade, over 220 million cases and 400 000 deaths were reported in 2018, with >90% of these occurring in the WHO African region. ^1^ More worryingly, a global stall in malaria control has been reported with a steady increase in malaria cases being observed between 2015 and 2018. ^1-2^ Caused by apicomplexan parasites of the genus *Plasmodium*, the most lethal form of human malaria is caused by *Plasmodium falciparum* which accounts for >99% of malaria cases and deaths in Sub-Saharan Africa. ^1^ However, human malaria caused by other *Plasmodium* spp. such as *P. vivax, P. ovale, P. malaria* and the zoonotic *P. knowlesi* remains a significant public health problem causing significant morbidity and economic impact in already poverty stricken communities. ^1^ The life cycle of malaria parasites comprises of multiple developmental stages between mosquito and mammalian hosts. Antimalarial drugs, which form principle components of malaria control programs, target the parasite at different life cycle stages, mostly the proliferating trophozoites and schizont stages during the intraerythrocytic development cycle of the parasites which are associated with most of the disease pathology. Artemisinins (ARTs) in ART combination therapies are the current front line drugs in malaria treatment. ^1^ They display fast and potent activity against virtually all blood stages of the parasites, as well as gametocytes that mediate transmission to mosquito vectors. ^3-4^ Indeed, such is the effectiveness of ARTs, that recent gains in malaria control have been partly attributed to ART combination therapies. ^2, 4^ Unfortunately, *P. falciparum* (PF) resistance to ARTs has emerged in the Southeast Asia greater Mekong region and is characterised by point mutations in the Kelch13 propeller domain that associate with decreased parasite clearance rates in clinical phenotypes. ^1, 4-5^

ARTs are sesquiterpene lactones derived from the Chinese herb *Artemisia annua*. Central to the activity of ARTs is the activation of the core endoperoxide bridge by haem which triggers the production of carbon centred radicals which in turn alkylate multiple and random downstream parasite targets. ^6-7^ The actual events leading to ART mediated parasite death remain elusive as well as disputed. However, a promiscuous targeting of several parasite proteins by the ART generated radicals is widely accepted. ^8-9^ The ART resistance-associated mutations lie in the beta propeller domain of the Kelch13 protein in PF. ^10^ Recent work on the biological function and consequences of these Kelch13 mutations has revealed that Kelch13 localises to the parasite cytoplasmic periphery in cellular compartments called cytostomes and plays a role in haemoglobin endocytosis. ART resistance-associated mutations in Kelch13 lead to reduced abundance of this protein leading to impaired haemoglobin trafficking which lessens ART activation hence promoting parasite survival. ^11-12^In addition, ART induced pleiotropic targeting is also known to activate ER stress and the unfolded protein response (UPR) which allow parasites to survive drug assault by rapidly turning over damaged proteins while employing cell repair mechanisms. ^6-7, 13^ ART resistant parasites (Kelch13 mutants) are indeed associated with an upregulation of genes involved in these cellular stress response pathways. ^14^ Meanwhile, parallel functional and localisation studies have also revealed that Kelch13 co-localises with multiple UPR components, proteins specific to the ER and mitochondria as well as intracellular vesicular trafficking Rab GTPases. ^15-16^ Central to the activity of the UPR is the ubiquitin proteasome system (UPS), a conserved eukaryotic pathway that plays a role in protein homeostasis by degrading unfolded proteins. Under ART pressure, activity of the UPS is more upregulated in Kelch13 mutant parasites compared to wild type while UPS inhibitors have been shown to synergize ART action suggesting that this pathway could be selectively targeted to overcome ART resistance. ^17-18^ Of note, Kelch13 is also predicted to play additional roles as substrate adaptor for ubiquitin E3 ligases, crucial components of the UPS; ^7, 10^ while mutations in upstream components of the UPS (ubiquitin hydrolases or deubiquitinating enzymes) also modulate susceptibility to ARTs. ^19-21^ Chemotherapeutic targeting of the UPS has been successfully pursued in cancers ^22^ and is increasingly becoming attractive in malaria parasites, ^23^ even more so as potential combinatorial partners to ARTs to overcome resistance. ^17-18^

Here we report the activity of deubiquitinating enzyme (DUBs) inhibitors in both rodent and human malaria parasites. DUBs are proteases that cleave ubiquitin residues from conjugated substrate proteins in the UPS pathway. UPS targeting of proteins is initiated by ubiquitin (Ub) tagging of substrates which marks them either for specific cellular signal transduction processes like DNA repair and cell cycle progression or subsequent degradation by the 20s proteasome. ^24^ Ub tagging is mediated by three sequential enzymes: E1, an activating enzyme; E2, a conjugating enzyme and E3, a Ub ligase for substrate specificity. The activity of these enzymes results in polyubiquitination of substrate proteins which signals for their degradation at the 20s proteasome complex depending on the number of Ub residues. DUBs reverse the activity of these downstream UPS enzymes by removing Ub from the conjugated substrates which results in diverse protein fates and cellular outcomes among which include; regulation of protein half-life, cell growth, differentiation, transcription; rescue of mis-tagged proteins as well as oncogenic and neuronal disease signalling. ^25^ Over 100 DUBs have been identified in humans and they classify into five major families: Ub C-terminal hydrolases (UCHs), Ub specific proteases (USPs), ovarian tumour proteases (OTUs), josephins and JAMM/MPN/MOV34. ^25^ In malaria parasites, up to 30 DUBs have been predicted across five *Plasmodium* species (PF, *P. vivax, P. berghei* (PB), *P. chabaudi, P. yoelii*); even though their functions remain to be fully explored. ^26-27^ Nevertheless, *Plasmodium* DUBs seem to have intrinsic protease activity, are significantly divergent and their human orthologues are known to be important regulators of cellular pathway which makes them suitable and potential drug targets. ^28^ The role of DUBs in mediating susceptibility to standard drugs like ARTs, the diversity in the classes of DUBs and the predicted repertoire in malaria parasites would also mean an expanded chemical space for drug discovery, potential inhibitor combination for different classes as well as using DUB inhibitor combinations to overcome ART resistance. Herein, using generic mammalian DUB inhibitors that have been used as exploratory research tools as well as in clinical trials, we show that DUB inhibitors do possess *in vitro* and *in vivo* inhibitory activities against malaria parasites across two diverged *Plasmodium* species. We demonstrate that different classes of DUB inhibitors can be combined to provide greater killing efficacy as well as enhance the potency of ARTs both *in vitro* and *in vivo*. Our data demonstrate that DUB inhibition can be exploited to overcome ART resistance with similar potency as first generation proteasome inhibitors. Furthermore, inhibition of both the UPS and DUBs can be combined to further improve the potency of ARTs and negate ART resistance. These findings have the potential to be applied to the treatment of all human malaria.

## Results

### *In vitro* activity of DUB inhibitors in malaria parasites

To assay for *in vitro* activity of DUB inhibitors in malaria parasites, short term PB culture assays and PF Sybergreen I^®^ culture assays were employed. The PB 820 and PF 3D7 lines were initially screened to determine susceptibility to inhibitors and antimalarials with known activity in malaria parasites; ART, dihydroartemisinin (DHA), chloroquine (CQ) and epoxomicin (20s proteasome inhibitor). The half-inhibitory concentrations (IC_50_) obtained for epoxomicin, DHA, ART and CQ in both the 820 and 3D7 lines (Table 1) were all in agreement with previously published IC_50_ values in both *Plasmodium* species. ^29-32^ Next, we screened seven DUB inhibitors (Table 1) in both the 820 and 3D7 line to characterise their inhibitory activity during the intraerythrocytic stages of malaria parasites. The selected compounds are DUB inhibitors being currently pursued as promising anticancer agents (Table 1) that also offered a broad coverage targeting of the 5 classes of DUBs. As shown in Table 1, activity was observed for six of the seven DUBs tested in the 820 and 3D7 lines. The activity of USP acting DUB inhibitors; b-AP15, P5091 and NSC632839 corresponds with the reported *in vitro* IC_50_s of the compounds screened in cancer cell lines ^33-35^. b-AP15 IC_50_ also compared to previously reported IC_50S_ of 1.54± 0.7 µM and 1.10 ± 0.4 μM in PF CQ sensitive (3D7) and resistant (Dd2) lines respectively. ^36^ Growth inhibition was also observed for broad spectrum DUB inhibitors; PR-619 and 1,10 phenanthroline, as well as a partially selective DUB inhibitor, WP1130 (Table 1). These data suggest that DUBs are potentially essential enzymes in *Plasmodium*, and they could be pursued as potential antimalarial drug targets. Indeed, a manual curation of up to 17 of the predicted DUBs in malaria parasites ^26-27^ shows that a majority of these (∼70%, 12 of 17) are essential in either PF and PB or both (Supplementary Table 1) based on previous functional studies for selected DUBs ^37-38^ or recent genome wide gene knockout screens. ^39-40^ Strikingly, no growth inhibition was observed for TCID (IC_50_ >100 µM), a UCH-L3 inhibitor, in both the 820 and 3D7 lines (Table 1, Supplementary Figure 1A, 1B). Among the well characterised DUBs in malaria parasites is PF UCH-L3 (PfUCH-L3, PF3D7_1460400) which was identified by activity based chemical profiling and has been shown to retain core deubiquitinating activity. ^41^ Structural and functional characterisation of PfUCH-L3 has also shown that this enzyme is essential for parasite survival (Supplementary Table 1). ^38^ Meanwhile, in our screen, TCID, a highly selective mammalian UCH-L3 inhibitor with an IC_50_ of 0.6 µM in mammalian cancer cell lines, ^42^ displayed no activity in both the 820 and 3D7 lines (Table 1, Supplementary Figure 1A, 1B). To possibly address this (unexpected) lack of activity, we performed a phylogenetic analysis of *Plasmodium*, human and mouse UCH-L3 based on predicted protein sequences to infer their similarities which might explain the observed lack of anti-plasmodial activity of TCID. A distinct evolutionary divergence of this enzyme was observed between human, mouse and the most similar *Plasmodium* homologues (PBANKA_ 1324100/PF3D7_1460400) which whilst annotated as UCH-L3 shares only 33% predicted protein sequence identity with the human UCH-L3 (Supplementary Figure 1C, D). Structurally, human UCH-L3 and PfUCH-L3 have similar modes of Ub recognition and binding. However, the PfUCH-L3 Ub binding groove is structurally different from the human UCH-L3 at atomic bonding level and possesses non-conserved amino acid residues. ^38^ This lack of complete identity across active sites would perhaps further explain the observed inactivity of TCID in both PF and PB.

**Table 1:** *In vitro* activity of DUB inhibitors in rodent and human malaria parasites. IC_50_ values and error bars are means and standard deviations from at least 3 independent repeats.

| Inhibitor | Predicted UPS target | IC<sub>50</sub> |  |
| --- | --- | --- | --- |
|  |  | PB 820 | PF 3D7 |
| Artemisinin | - | 17.23±0.4nM | 6.50±0.4nM |
| Dihydroartemisinin | - | 13.89±0.1nM | 6.23±0.34nM |
| Epoxomicin | 20s proteasome | 14.20±3.0nM | 11.12±0.23nM |
| PR-619 | broad spectrum DUB inhibitor<sup>a</sup> | 3.30±2.0μM | 2.41±0.5μM |
| P5091 | USP7 and USP47 DUBs<sup>b</sup> | 8.38±2.10μM | Not done |
| TCID | UCH-L3 and UCH-L1 DUBs<sup>c</sup> | >100μM | >100μM |
| WP1130 | UCH-L1, USP9X, USP14, UCH37 DUBs<sup>d</sup> | 1.19±1.0μM | 2.92±0.1μM |
| b-AP15 | USP14 and UCH-L5 DUBs<sup>e</sup> | 1.06±0.9μM | 1.55±0.1μM |
| NSC-632839 | USP2, USP7, SENP2 DUBs<sup>f</sup> | 27.97±0.8μM | Not done |
| 1,10 phenanthroline | Metalloproteases and JAMM isopeptidases<sup>g</sup> | 0.63±0.3μM | Not done |

a, <sup>64</sup> b, <sup>33</sup> c, <sup>42</sup> d, <sup>65</sup> e, <sup>34</sup> f, <sup>35</sup> g. <sup>66</sup>

### Different classes of DUB inhibitors can be combined to provide more effective blocking of malaria parasite growth *in vitro*

To explore interactions between DUB inhibitors, and their potential synergy, b-AP15, a highly selective USP14 inhibitor ^34^ and the relatively most potent inhibitor of parasite growth in both PF and PB, was tested in fixed ratios with broad-spectrum DUB inhibitors; PR-619 and WP1130. Combinations at fixed ratios of 5:0, 4:1, 3:2, 1:4 and 0:5 were serially diluted and incubated with parasite cultures of the 3D7 line from which parasite growth and IC_50_s were obtained. FIC_50_s and ∑FIC_50_s were calculated and isobologram interactions were plotted. A combination of b-AP15 and PR-619 is mostly additive with a mean ∑FIC_50_ of 0.753±0.23, (Figure 2A). Meanwhile, b-AP15 and WP1130 seemingly trends towards synergy with a mean ∑FIC_50_ of 0.653±0.23, (Figure 2B) even though the interaction remains overall additive. These data suggested that DUB inhibitors, as potential antimalarial drug candidates, can be used in combination to block parasite growth presumably by simultaneously targeting several different DUB enzymatic targets.

**Figure 1:**
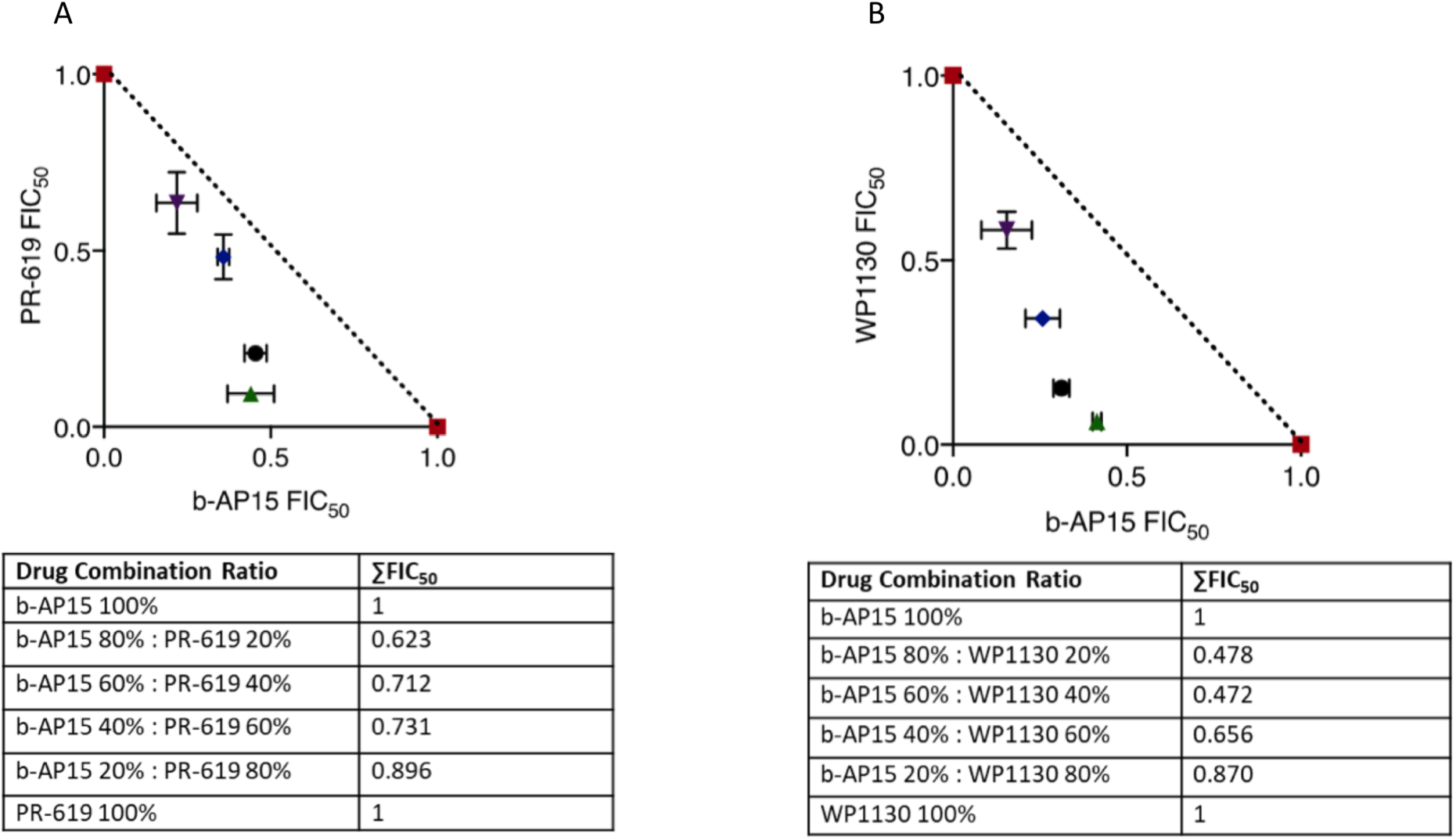
*In vitro* interaction of different classes of DUB inhibitors in malaria parasites. Isobologram interaction plots and ∑FIC_50_ values of interactions between DUB inhibitors in the PF 3D7 line. **A**. Interaction between b-AP15 and WP1130 and their raw ∑FIC_50_ values. **B**. Interaction between b-AP15 and PR-619 and their raw ∑FIC_50_ values. ∑FIC_50_ values, plotted FIC_50_s and error bars are means and standard deviations from three biological repeats.

**Figure 2:**
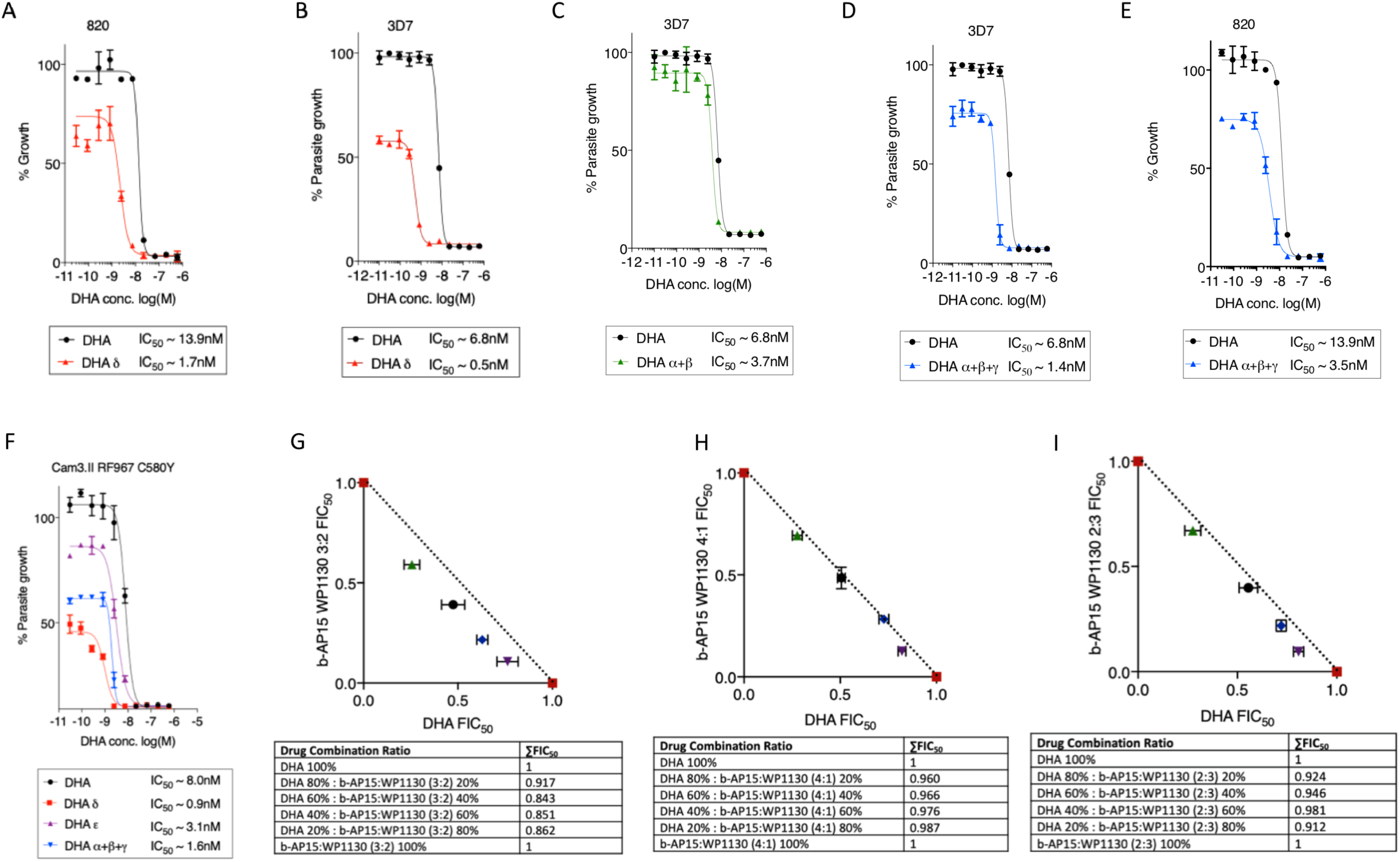
*In vitro* potentiation of DHA by DUB inhibitors. **A, B**. Dose response profiles and IC_50_ values of DHA in the presence of b-AP15 at IC_50_ equivalent concentration (DHA δ) in the PB 820 line (**A**) and 3D7 line (**B**). **C**. Dose response profiles and IC_50_ values of DHA in the presence of WP1130 and PR-619 at their respective half IC_50_s (DHA α+β) in the 3D7 line. **D, E**. Dose response profiles and IC_50_ values of DHA in combination with b-AP15, WP1130 and PR-619 at half IC_50_ (DHA α+β+γ) in the 3D7 (**D**) and 820 line **(E). F** Dose response profiles and IC_50_ values of DHA combined with b-AP15 and WP1130 at IC_50_ (DHA δ, DHA ε) or b-AP15, WP1130 and PR-619 at half IC_50_ (DHA α+β+γ) in ART resistant Kelch13 C580Y mutant line. Dose response curves were plotted in Graph pad prism 7. Error bars are standard deviations from 3 independent biological repeats. Isobologram plots of DHA in combination with b-AP15 and WP1130 at 3:2 (**G**), 1:4 (**H**) and 2:3 (**I**) ratios and their raw ∑FIC_50_ values. ∑FIC_50_ values, plotted FIC_50_s and error bars are means and standard deviations from three biological repeats.

### DUB inhibitors alone or in combination can potentiate DHA action in malaria parasites *in vitro*

In order to test the hypothesis that DUB inhibitors might have a similar effect of potentiating ART activity as 20s proteasome inhibitors, we investigated the effects of DUB inhibitors on the dose response profiles of DHA *in vitro* on wild type PB and PF growth as well as their potential to synergize DHA action in fixed ratio interaction assays. The most potent DUB inhibitor b-AP15 at equivalent IC_50_ concentration improved DHA action with up to ∼8-fold IC_50_ shift in wild type PB growth inhibition (Figure 2A) and up to 15-fold enhancement in the wild type PF growth inhibition (Figure 2B). The differences in potentiation between PB and PF could be due to the inherent reduced susceptibility of PB to ARTs. ^20, 43^ The enhancement of DHA action by b-AP15 was also almost similar to previously reported profiles with epoxomicin, a 20s proteasome inhibitor. ^17^ We have recently shown that experimental introduction of mutations in a DUB, UBP-1, mediated reduced susceptibility to ARTs in PB. ^20^ UBP-1 has a close human orthologue HAUSP/USP7 which is itself inhibited by P5091, a drug which in our *Plasmodium* screen was poorly potent with a relatively high micromolar IC_50_ (Table 1). Nevertheless, b-AP15 (a USP-14 inhibitor) potentiated DHA action to the same extent as in wild type ART-sensitive PB (9-11-fold) in two UBP1 mutant lines that have reduced susceptibility to ART (V2721F) or both ART and CQ (V2752F) (Supplementary Figure 2A & B). Therefore, ART (and potentially CQ) reduced susceptibility could be offset by a combinatorial drug administration approach involving DUB inhibitors through a targeted disruption of protein homeostasis most likely at the level of the UPS.

In an attempt to maximise DUB inhibitor combinations, which offered improved inhibition of parasite growth (Figure 1) as a strategy for simultaneously targeting several DUBs in the presence of DHA, we tested the effect of combining b-AP15, PR-619 and WP1130 on the dose response profile of DHA. WP1130 and PR-619 at IC_50_ concentration mildly potentiate DHA action with 1.8- and 1.4-fold improvements respectively (Supplementary Figure 3A, 3B). Meanwhile, a combination of b-AP15 and WP1130 at half IC_50_ mildly potentiated DHA action (∼2-fold, Figure 2C), while all three inhibitors (b-AP15, WP1130 and PR-619) at half IC_50_ improved DHA action up to 5-fold in the ART sensitive PF (Figure 2D) and PB (Figure 2E) as well as the ART resistant PF Kelch13 C580Y mutant lines (Figure 2F).

We carried out further isobologram interaction assays for DUB inhibitor ratio combinations in an attempt to achieve improved *in vitro* killing (Figure 1) in combination with DHA. Both b-AP15 and WP1130 were essentially additive when combined with DHA in isobologram interactions with ∑FIC_50_s of 0.967 and 1.013 respectively (Supplementary Figure 3C, 3D). However, when b-AP15 and WP1130 were mixed at a 3:2 molar concentration ratio as a cocktail and combined with DHA, a slight improvement in efficacy was observed with an ∑FIC_50_ of ∼0.868 (Figure 2G) compared with 0.972 at 1:4 b-AP15 WP1130 molar concentration ratios (Figure 2H) or 0.941 at 2:3 b-AP15 WP1130 molar concentration ratio (Figure 2I). These data would suggest that optimized ratios of (improved) DUB inhibitor combinations or other proteasome inhibitors might yet achieve synergy with DHA, which would be a prerequisite to simultaneously targeting multiple DUBs or parallel pathways/enzymes in the UPS in future antimalarial combination therapies.

### A combination of DUB and 20s proteasome inhibitor can synergize with DHA

An alternative approach to alleviating antimalarial resistance is combination therapies that target multiple points within known resistance mediating pathways and/or novel antimalarial drug pathways to prevent the emergence of or overcome resistance. Therefore, we explored a combination of an upstream DUB inhibitor (b-AP15) and a 20s proteasome inhibitor (epoxomicin) with DHA in fixed ratio isobologram interactions. Firstly, we tested epoxomicin in combination with DHA as well as b-AP15 and epoxomicin in fixed ratios against PF. Epoxomicin improved DHA action mildly with an ∑FIC_50_ of 0.881 (Figure 3A) which corresponds with previously reported profiles. ^17^ Interestingly, b-AP15 and epoxomicin as a combination alone was not an improved regimen with an ∑FIC_50_ of 1.162 (Figure 3B). This failure may result from a suppression mechanism where targeting the USP14 DUB upstream by b-AP15 (Figure 3D) would potentially counteract the activity of downstream 20s proteasome inhibitor and vice versa. ^44^ However, a 1:1 molar ratio of b-AP15 and epoxomicin when combined with DHA, an improved interaction with DHA (∑FIC_50_ of 0.614) was achieved (Figure 3C) than by either of the drugs alone (Figure 3A, Supplementary Figure 3C). This illustrates that targeting the UPS at several points with the optimized inhibitor concentrations can significantly improve DHA efficacy.

**Figure 3:**
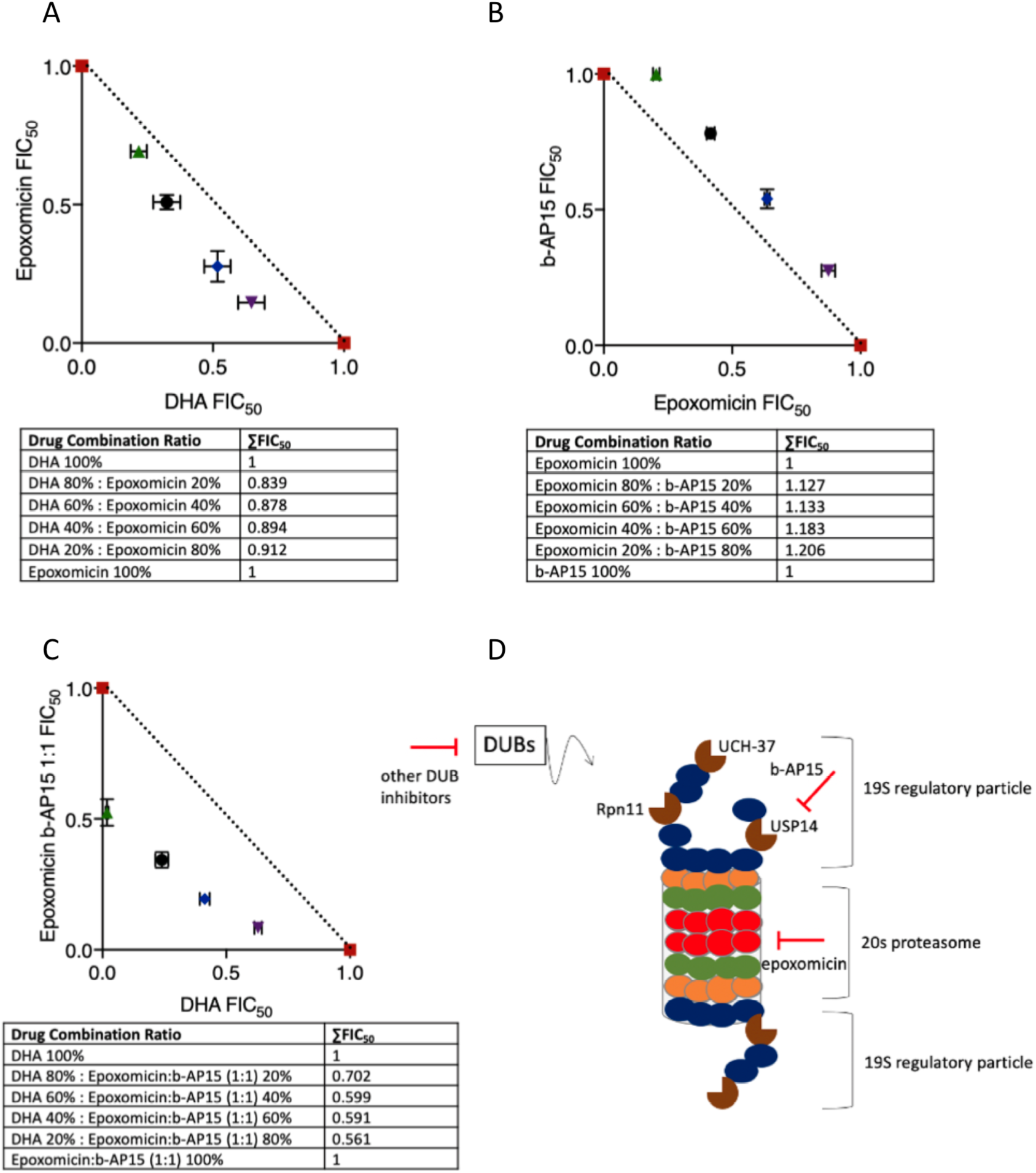
A combination of DUB and 20s proteasome inhibitor improves synergy with DHA. **A-C**. Isobologram interaction between epoxomicin and DHA **(A)**, b-AP15 and epoxomicin **(B)** and a mixture of b-AP15 and epoxomicin at 1:1 molar concentration ratio in combination with DHA **(C)**. ∑FIC_50_ values, plotted FIC_50_s and error bars are means and standard deviations from three biological repeats. **D**. Illustrated figure of the UPS indicating positional scope of USP14 and 20s units of the UPS and the inhibitor targets.

### Pre-incubation of malaria parasites with UPS inhibitors efficiently mediates DHA potentiation

A further way to combat drug resistance in malaria, which is being explored with antibiotics ^45^ and has been the case with cancer neo-adjuvant therapies, would be to pre-expose parasites to lethal or sub-lethal doses of inhibitors that target the resistance pathways before the main treatment course. A targeted inhibition of the resistance conferring pathways might then in turn improve the activity of any downstream main treatment drug. Therefore, we investigated the effect of pre-exposing malaria parasites to DUB or 20s proteasome inhibitors on the short time exposure dose response profiles to DHA in both PB and PF. The PB 507 line, which expresses a green fluorescent protein (GFP) constitutively, was used to monitor GFP intensity across the life cycle after exposure to serial concentrations of DHA for 3 hours, administration of which followed prior exposure of the parasites (1.5 hour old rings) for 3 hours to IC_50_ concentrations of b-AP15. Quantification of the GFP fluorescent signal expressed from a constitutive promoter in PB would allow us to investigate the global dynamics of protein homeostasis, recycling, unfolding and or damage which occurs in the parasites upon exposure to DHA and or UPS inhibitors. Monitoring of GFP intensity at 6, 18 and 24 hours revealed that b-AP15 pre-exposure enhances the potency of DHA as indicated by significant abrogation of GFP intensity at all the time points (Figure 4A). Additional administration of b-AP15 after DHA incubation further abrogates GFP intensity illustrating that b-AP15 compromises UPS activity in tandem with DHA, which would make them suitable partner drugs. In the PF 3D7 line, pre-incubation of ∼0-3 hour old rings with b-AP15 at IC_50_ or half IC_50_ for 3 hours followed by DHA treatment for 4 hours markedly impacts parasite viability (5 and 1.6 fold respectively) compared to DMSO exposed parasites, while pre-exposing the parasites to b-AP15 at 4x IC_50_ is almost entirely lethal to the parasites (Figure 4B). Meanwhile, pre-exposure of 3D7 or an ART resistant Kelch13 C580Y line to epoxomicin at IC_50_ or 0.2x IC_50_ followed by DHA also significantly impacted parasite viability (∼4.6 and ∼1.4 fold respectively) as compared to DMSO (Figure 4C, 4D). Remarkably, in both the 3D7 and ART resistant Kelch13 C580Y lines, a combination of b-AP15 and epoxomicin at half IC_50_ achieved better potency with DHA (18 and 33-fold respectively) compared to either of the drugs alone at IC_50_ (Figure 4B, 4C, 4D) further illustrating that targeting multiple UPS components (Figure 3C) could be a flexible approach to overcoming ART resistance.

**Figure 4:**
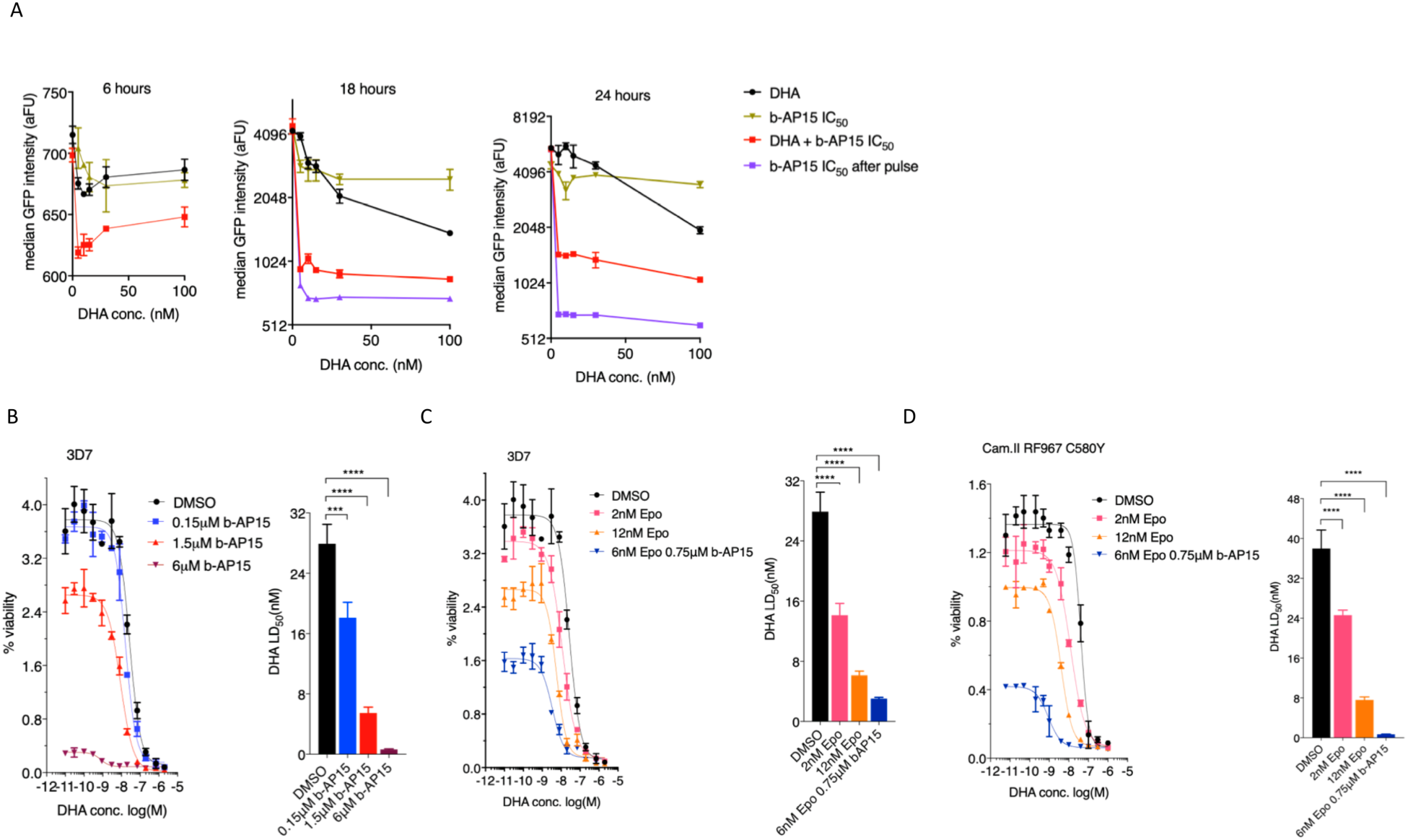
pre-exposure of malaria parasites to UPS inhibitors alone or in combination enhances DHA action. **A** pre-treatment of the PB 507 line (1.5 hours old rings) with b-AP15 at IC_50_ (1.5µM) for 3 hours followed by a wash and then DHA for another 3 hours. Median GFP intensity quantified by flow cytometry at 6 hours, 18hours and 24 hours. b-AP15 at IC_50_ readded after DHA wash off in one experimental condition (green plot) while b-AP15 alone used as an additional control. Results are representative of three independent experiments. **B**. DHA dose response viability plots and lethal dose (LD_50_) comparisons at 66 hours after pre-exposure of 0-3 hours old rings of the 3D7 line to DMSO (0.1%) or b-AP15 at half IC_50_ (0.75µM), IC_50_ (1.5µM) or 4X IC_50_ (6µM) followed by DHA for 4 hours. **C, D**. DHA dose response viability plots and lethal dose (LD_50_) comparisons at 66 hours after pre-exposure of 0-3 hours old rings of the 3D7 line (**C**) and ART resistant Kelch-13 C580Y line (**D**) to DMSO (0.1%) or epoxomicin at 0.2x IC_50_ (2nM), IC_50_ (12nM) or a combination of b-AP15 and epoxomicin at half IC_50_ followed by DHA for 4 hours. Data from three independent experimental repeats. Significant differences between the conditions were calculated using one-way ANOVA alongside the Dunnet’s multiple comparison test. Significance is indicated with asterisks; ****p <0.0001.

### b-AP15 fails to block parasite growth but potentiates ART action *in vivo*

We next investigated the ability of b-AP15 to block parasite growth *in vivo* and potentially enhance ART action. An analogue of b-AP15 (itself a lead first generation DUB inhibitor), VLX1570 entered clinical trials for the treatment of multiple myeloma (Wang et al., 2016), despite being later terminated due to dose ascending toxicities (NCT02372240). b-AP15 has strong antiproliferative effects in human cancer cell lines and has displayed significant antitumor activity at 5mg/kg in *in vivo* mouse models without any side effects. ^34^ However, in a Peters’ 4 day suppressive test, b-AP15 fails to clear PB parasites *in vivo* at both 1mg/kg and 5mg/kg with only minor reductions in parasite burdens on day 4 and 5 post treatment at the latter dose which corresponds to ∼70% parasite suppression on day 4 (Figure 5A, 5B, 5C). Contrary to the previous reported safety profiles of b-AP15, ^34^ mice (Theiler’ s Original) treated with 5mg/kg b-AP15 started to develop toxicity signs as demonstrated by significant weight loss on day 4 and 5 post-treatment. Further treatments at 5mg/kg or higher doses were thus not pursued. To investigate the ability of b-AP15 to potentiate ART action *in vivo*, b-AP15 was administered at 1mg/kg (a safe dose that did not have any effect on parasite growth alone, Figure 5A) in combination with ART at 5mg/kg and 10mg/kg in established mice infections at a parasitaemia of 2-2.5% for three consecutive days. A combination of ART (5mg/kg) and b-AP15 (1mg/kg) did not have any significant parasite reduction as compared to ART (5mg/kg) alone, while ART at 20mg/kg cleared the parasites after three consecutive doses as expected (Figure 5D). However, a combination of ART (10mg/kg) and b-AP15 (1mg/kg) significantly abrogated parasite burden as compared to ART (10mg/kg) alone to the same extent as ART at 20mg/kg (Figure 5E). These data further showed that b-AP15 can enhance ART action *in vivo*, to a similar extent as observed *in vitro*.

**Figure 5:**
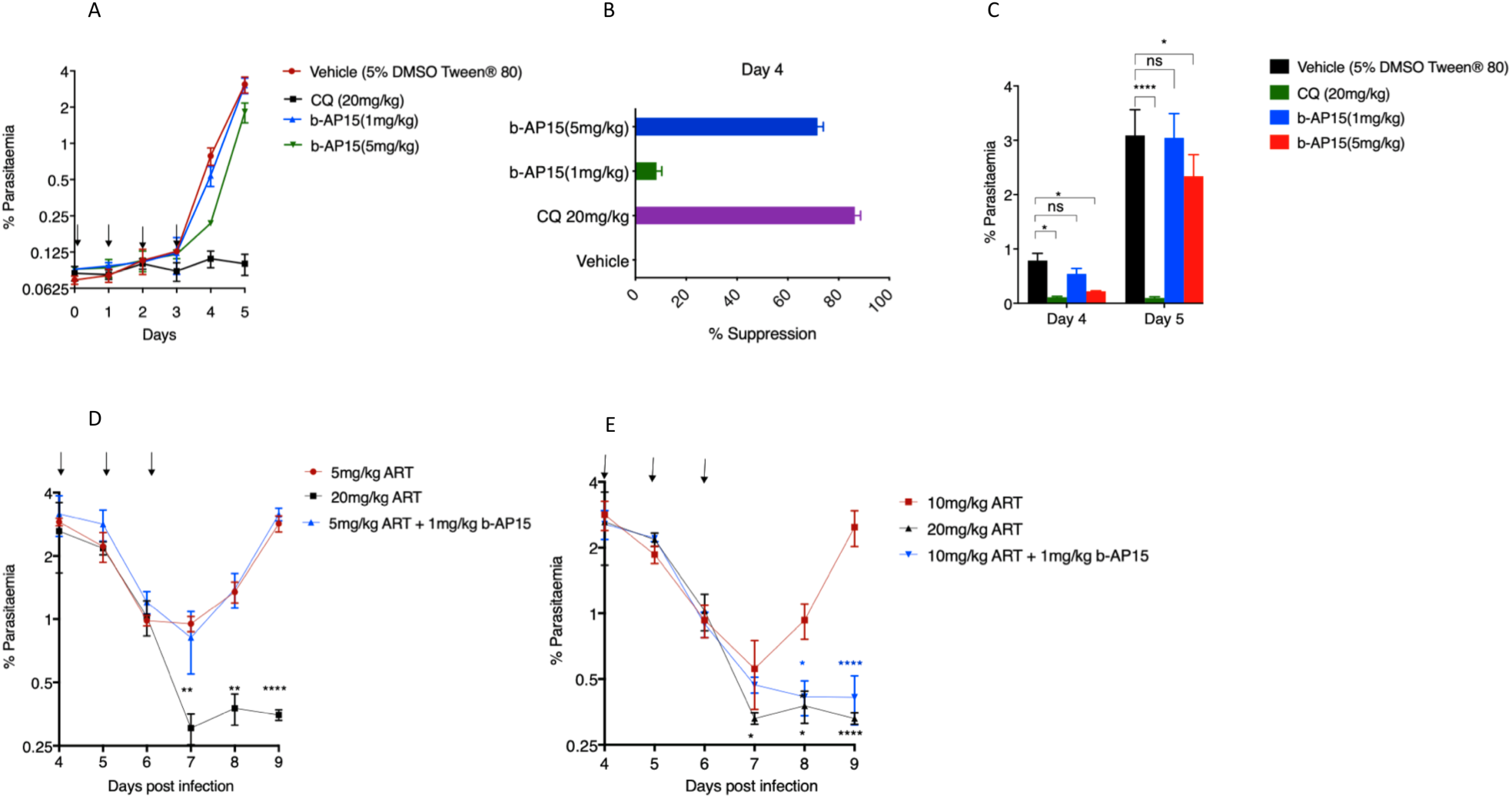
*In vivo* activity of b-AP15 alone and or in combination with ART. **A** Mice (4 groups of 3 mice each) were infected with 10^5^ parasites on day 1 and treated with indicated drug doses ∼1 hour post infection for four consecutive days (indicated by arrows). Parasitaemia was monitored daily by flow cytometry and analysis of Giemsa stained smears. **B, C**. Percentage suppressions on day 4 (**B**) and bar of parasitaemias on day 4 and day 5 (**C**). **D, E**. Combination of ART and b-AP15 in established mouse infections. ART at 5mg/kg (**D**) or 10mg/kg (**E**) combined with b-AP15 (1mg/kg) administered in established mice infections at a parasitaemia of 2-2.5% for three consecutive days (indicated by arrows). Parasitaemia was monitored daily. ART at 20mg/kg was used as a curative control. Significant differences were calculated using one-way ANOVA alongside the Dunnet’s multiple comparison test. Significance is indicated with asterisks; *p <0.05, **p <0.01, ***p <0.001, ****p <0.0001.

## Discussion

With the increasing incidence of resistance to (even combinations of) antimalarial drugs by PF and the lack of rapidly amenable drug discovery programs for related *Plasmodium* spp. such as *P. vivax*, pipelines to develop new antimalarial drugs to treat the disease as well as improve the activity of current antimalarials and tackle resistance are urgently needed. Here, we report *in vitro* and *in vivo* activity of a class of compounds targeting the parasite upstream UPS component (DUBs) in PF and PB. Antimalarial drugs are typically discovered for their activity against PF *in vitro*. Lead compounds from PF *in vitro* screens are evaluated for *in vivo* efficacy using rodent malaria parasites which have been for a long time, crucial components of these drug discovery programs. ^46^ PB is the most commonly used rodent model (in what is called the Peters’ four-day suppressive test) and the development of methods that allow assessment of both *in vitro* drug sensitivity and *in vivo* efficacy in this model, ^47^ as we demonstrate in this study, permits easy comparisons with PF *in vitro* efficacy data. Moreover, this provides crucial *in vitro* bridging information on whether potential drug efficacy discrepancies between PF *in vitro* and PB *in vivo* are due to pharmacokinetics of the drug or intrinsic differences in drug sensitivity between the *Plasmodium* spp. As a species of *Plasmodium* that is well diverged from both PF and other human-infectious *Plasmodium*, PB drug efficacy assessment also offers a useful comparative for other non-PF human causing *Plasmodium* spp. as chemical entities that display PF inhibitory activity *in vitro* and PB inhibitory activity *in vitro* and *in vivo* are also likely to be active against other (human infectious) *Plasmodium* species.

Herein, activity is reported for six DUB inhibitors covering most of the DUB enzyme families and include b-AP15, P5091 and NSC632839 which specifically target USPs that all displayed antimalarial activity against both rodent and human malaria parasites *in vitro*. USPs are the largest family of DUBs comprising of up to 56 individual enzymes in humans. ^48^ However, since less is known of USPs in malaria parasites, with their current assignations largely based on in silico predictions, ^26-27^ the precise targets of these drugs remain largely obscure. Human USP14 has been demonstrated to be the target of b-AP15 ^34^ and its PF orthologue PfUSP14 (PF3D7_0527200) has been recently characterised and shown to bind the parasite 20s proteasome. ^36^ Moreover, purified PfUSP14 cleaves di-ubiquitin bonds in intact polyubiquitin chains illustrating functional identity of this *Plasmodium* DUB with its human counterpart. ^36^ This provides evidence that PfUSP14 may be specifically essential in parasite proliferation during the asexual blood cycle which was supported by a whole genome piggyBac saturation mutagenesis screen in which PfUSP14 was shown to be refractory to deletion (Supplementary Table 1). ^39^ Our data also support this in both PF and PB despite the PB counterpart (PBANKA_1242000) appearing to be dispensable in a recombinase mediated genetic screen. ^40^ The differences in essentiality could be due to functional differences between the two *Plasmodium* spp. USP14s. as they seem to share only ∼62% sequence identity (Supplementary Figure 4). The activity of b-AP15 in both PF and PB however, at almost equivalent potencies, could thus be suggestive of possible suitable compensatory effects from other DUBs upon deletion in PB which is not sufficiently compensated for when an inhibitor is used. b-AP15 may also target other DUB (or possess off target) activities in *Plasmodium* as the inhibition of purified PfUSP14 by b-AP15 is less potent than its overall parasite killing potency. ^36^ Nevertheless, the observed structural difference between human USP14 and PfUSP14 at the core catalytic domain, its possible essentiality and the activity of b-AP15 in both PF and PB *in vitro* suggests that PfUSP14 can be selectively targeted throughout the *Plasmodium* genus. ^36^ Furthermore, the observed activity of other USP inhibitors, P5091 and NSC632839 in this study suggests that their targets are essential (Supplementary Table 1) during the asexual proliferation stages of malaria parasites and can serve as useful chemical leads for more potent antimalarial discovery. More importantly, b-AP15 possesses antiparasitic activity *in vivo* achieving up to 70% parasite suppression of PB at the highest concentrations that have been tested in cancer models. ^34^ Malaria parasites have been shown to rapidly replenish proteasomes in the presence of sub-lethal doses of proteasome inhibitors ^49^ which would possibly explain the observed inability of b-AP15 to completely block parasite growth at this concentration as compared to control antimalarial drugs. Whilst promising, we noted issues with the reported safety profiles of b-AP15 at 5mg/kg ^34^ where mice significantly lost weight after 4 consecutive doses. This effect could be due to the combination of a chemical inhibitor and parasite challenge making the mice more susceptible to toxic effects of b-AP15, a phenomenon which has been previously reported with carfilzomib, a 20s proteasome inhibitor. ^49^ Meanwhile, the *in vitro* activity of broad-spectrum DUB inhibitors, PR-619 and WP1130 as well as a zinc chelating metalloprotease inhibitor (1, 10 phenanthroline) further alludes to the promise of DUBs as drug targets in malaria parasites.

A further striking finding was the inactivity of TCID (a UCH-L3 inhibitor) in both rodent and human malaria parasites. PfUCH-L3 has been well characterised in malaria parasites and has been shown to retain core deubiquitinating activity. ^41^ Moreover, disruption of PfUCH-L3 by experimentally replacing the native enzyme with a catalytically dead form was shown to be lethal to the parasite. ^38^ The inactivity of TCID in both rodent and human malaria parasites reported here is therefore suggestive of striking differences between mammalian and *Plasmodium* UCH-L3s. Our sequence analysis demonstrated that PfUCH-L3 shares ∼33% sequence identity with human UCH-L3 consistent with previous structural and molecular docking comparisons of PfUCH-L3 and human UCH-L3 which also revealed significant differences between the enzymes especially at the ubiquitin binding groove. ^38^ This makes PfUCH-L3 an even more attractive drug target for ultra-selectivity as it is also known to possess denedylating activities which are absent in mammalian UCH-L3s. ^41^

Targeting the *Plasmodium* UPS is an emerging interventional point, not just as a potential drug target, but now also to curb emerging ART resistance. 20s proteasome inhibitors have been shown to enhance ART action in both ART sensitive and resistant lines. ^17-18^ Our data in this study also show that upstream targeting of the UPS by some but by no means all DUB inhibitors can potentiate and enhance ART action in certain cases to a similar extent as 20s proteasome inhibitors. ARTs act by targeting several (possibly random) parasite proteins upon activation ^8-9^ which necessitates, among other things, an upregulated UPS mediated stress response which rapidly recycles and clears damaged proteins henceforth promoting survival in ART resistant parasites. ^6, 13, 17^ As with 20s proteasome inhibitors, ^17-18^ inhibition of parasite UPS by targeting single or multiple DUBs simultaneously potentiates ART or DHA action. Inhibition of parasite UPS by b-AP15, for example, would prevent the normal protein homeostasis flux through the UPS, boosting the activity of pleiotropic ARTs by blocking the parasite stress and recovery system. Indeed, despite DHA being only additive in our isobole study with b-AP15, sublethal concentrations of b-AP15 can boost DHA activity up to 15-fold. This boost is further enhanced when 2-3 DUB inhibitors at sub-lethal concentrations are combined as they improve DHA activity more than either inhibitors alone. This suggests that carefully titrated use of current DUB inhibitors in isolation, or simultaneously in mixtures may be a means to overcome ART resistance and the rodent model deployed here could be useful tool to optimise drug dosages. Indeed, recent findings have shown that accumulation of polyubiquitinated proteins in malaria parasites either by DUB or 20s proteasome inhibition is critical in activating the stress responses and contributes to DHA lethality in malaria parasites. ^13^ The observed increase in ART efficacy when combined with DUB inhibitors which is of a similar level to that achieved by inhibition of the proteasome by epoxomicin *in vitro* and Carfilzomib *in vivo* ^17^ further alludes to the potential of DUB inhibitors for achieving similar attributes in malaria parasites.

Indeed, whilst useful as independent potential antimalarial agents, DUB inhibitors show potential for partnership and this study demonstrated that different classes of DUBs can be targeted simultaneously to achieve better parasite killing while potentially minimising the resistance emergence window. More importantly, low and safe doses of b-AP15 with no effect on parasite growth alone significantly potentiated sub-curative dose of ART to almost curative levels *in vivo* providing a proof of concept that DUB inhibitors can enhance the activity of ARTs both *in vitro* and *in vivo* making them potential adjunct drugs to enhance ART action and tackle resistance. Similarly, other potential radical ways of overcoming resistance in malaria parasites would be combining drugs with different mode of actions in complex combinations or using multiple (different) first line combinational therapies at once to raise the probability barrier of developing resistance by simultaneously targeting several pathways. ^50^ Our data exemplify this concept, as for example when b-AP15 and epoxomicin are combined in a fixed ratio isobole analysis, their appears to be no interaction or possibly even an antagonistic effect. This observation would be symptomatic of an antagonistic suppression mechanism where the activity of two inhibitors in the same pathway upstream or downstream negatively feeds back to the activity of the other leading to counteractive effects. However, when b-AP15 and epoxomicin are mixed in equal concentration ratios and combined with DHA, their overall activity achieves a better efficacy with DHA than either of the inhibitors alone. The optimal simultaneous exposure of the parasite UPS to DUBs and 20s proteasome inhibitors could thus act as an additional opportunity to overcome resistance to ARTs if the parasites would acquire resistance mutations to either of the UPS inhibitors. This has indeed been recently illustrated where combined inhibition of the parasite β2 and β5 subunits of the parasites UPS has been shown to strongly synergize DHA activity. ^51^

In conclusion, our work confirms DUBs as potential druggable candidates in malaria parasites. Drug discovery programs take a long time, with for example a minimum of five years required to take a lead compound to a clinical candidate in malaria. ^52-53^ The emergent resistance to ACTs, a paucity in the number of antimalarial drugs in the developmental pipeline and a lack of scalable pipelines for drug discovery in other human malaria parasites such as *P. vivax and P. ovale*, ^53^ all necessitates both radical as well as alternative approaches to identify new drugs and drug targets. As DUBs are already being actively explored as anticancer agents with candidate inhibitors already entering clinical trials, ^54^ antimalarial drug discovery programs could take advantage to structurally improve or re-purpose such entities not just as potential drug targets in malaria, but also as combinational partners to ARTs to overcome the spectre of resistance.

## Materials and methods

### Parasite lines

Experiments in PB were carried out in an 820 line that expresses green fluorescent protein (GFP) and red fluorescent protein (RFP) in male and female gametocytes respectively, and a 507 line that constitutively expresses GFP under the control of the Pbeef1αa promoter. Generation and characterisation of the 820 and 507 lines has been previously described. ^55-56^ Growth inhibitory experiments in PF were performed in the CQ and ART sensitive 3D7 line and the ART resistant Cambodian Kelch13 C580Y mutant line (a kind gift from D. Fidock).

### Drugs and inhibitors

DHA (Selleckchem) was prepared at 1mM stock concentration in 100% DMSO and diluted to working concentration in complete (PF) or schizont media (PB). ART (Sigma) and Epoxomicin (Sigma) were dissolved in 100% DMSO to stock concentrations of 100 µM and 90µM respectively and diluted in complete culture media or schizont culture media to their respective working concentrations. CQ diphosphate (Sigma) was dissolved to stock concentration of 10 mM in 1X phosphate buffered saline (PBS) and diluted to working concentration in complete or schizont culture media. Seven different classes of DUB inhibitors (Table 1) were screened and were all obtained from Focus Biomolecules except for 1, 10 phenanthroline which was obtained from BPS biosciences. Stocks of DUB inhibitors were prepared at 10 mM in 100% DMSO and diluted in complete or schizont media to working concentrations. Testing concentrations ranged from 2000-0.01nM for epoxomicin, DHA, ART and CQ and 100-0.002µM for DUB inhibitors. All DUB inhibitors were supplied at a purity grade of >97% (Supplementary Table 2) and further analysed for chemical integrity on a High-Performance Liquid Chromatography (HPLC) platform (Supplementary Table 3, Supplementary Figure 5) as detailed below.

### HPLC analysis of DUB inhibitors

HPLC solvents were purchased from standard suppliers and used without additional purification. DUB inhibitors were analysed on a Shimadzu reverse-phase HPLC (RP-HPLC) system equipped with Shimadzu LC-20AT pumps, a SIL-20A auto sampler and a SPD-20A UV-vis detector (monitoring at 254 nm) using a Phenomenex, Aeris, 5 µm, peptide XB-C18, 150 × 4.6 mm column at a flow rate of 1 mL/min. RP-HPLC gradients were run using a solvent system consisting of solution A (H_2_O + 0.1% trifluoroacetic acid) and B (acetonitrile + 0.1% trifluoroacetic acid). Further gradient analyses were run from 0% to 100% using solution B over 20 minutes. Analytical RP-HPLC data was reported as column retention time in minutes. Percentage purity was quantified by percentage peak area in relation to main peak.

### PB animal *infections*

PB parasites were maintained in female Theiler’s Original (TO) mice (Envigo) weighing between 25-30g. Parasite infections were established either by IP of ∼200µl of cryopreserved parasite stocks or intravenous injections (IVs) of purified schizonts. For infections from a donor infected mouse (mechanical passage), 5-30µl of infected blood was diluted in phosphate buffered saline (PBS) followed by injections of 100-200µl by IP. Since PB preferentially invades reticulocytes, ^57^ mice were pre-treated with 100µl of phenylhydrazine at 12.5mg/ml in physiological saline 2 days before the infections to induce reticulocytosis for some experiments. Routine monitoring of parasitaemia in infected mice was done by monitoring methanol fixed thin blood smears stained in Giemsa (Sigma) or flow cytometry analysis of infected blood stained with Hoescht 33342 (Invitrogen). Blood from infected mice was collected by cardiac puncture under terminal anaesthesia. All animal work was approved by the University of Glasgow’s Animal Welfare and Ethical Review Body and by the UK’s Home Office (PPL 60/4443) and carried out by appropriately licenced individuals. The animal care and use protocol complied with the UK Animals (Scientific Procedures) Act 1986 as amended in 2012 and with European Directive 2010/63/EU on the Protection of Animals Used for Scientific Purposes.

### PB *in vitro* culture and drug susceptibility assays

For *in vitro* maintenance of PB, cultures were maintained for one developmental cycle using a standardised schizont culture media containing RPMI1640 with 25mM hypoxanthine, 10mM sodium bicarbonate, 20 % fetal calf serum, 100U/ml Penicillin and 100μg/ml streptomycin. Culture flasks were gassed for 30 seconds with a special gas mix of 5% CO2, 5% O2, 90% N2 and incubated for 22-24 hours at 37°C with gentle shaking, conditions that allow for development of ring stage parasites to mature schizonts. Drug assays to determine *in vitro* growth inhibition during the intraerythrocytic stage were performed in these standard short-term cultures as previously described. ^29-30^ Briefly, 1 ml of infected blood with a non-synchronous parasitaemia of 3-5% was collected from an infected mouse and cultured for 22-24 hours in 120 ml of schizont culture media. Schizonts were enriched from the cultures by Nycodenz density flotation as previously described ^58^ followed by immediate injection into a tail vein of a naive mouse. Upon IV injection of schizonts, they immediately rupture with resulting merozoites invading new red blood cells within minutes to obtain synchronous *in vivo* infection containing >90% rings and a parasitaemia of 1-2%. Blood was collected from the infected mice 2 hours post injection and mixed with serially diluted drugs in schizont culture media in 96 well plates at a final haematocrit of 0.5% in a 200µl well volume. Plates were gassed and incubated overnight at 37° C. After 22-24 hours of incubation, schizont maturation was analysed by flow cytometry after staining the infected cells with DNA dye Hoechst-33258. Schizonts were gated and quantified based on fluorescence intensity on a BD FACSCelesta or a BD LSR Fortessa (BD Biosciences, USA). To determine growth inhibitions and calculate IC_50_, quantified schizonts in no drug controls were set to correspond to 100% with subsequent growth percentages in presence of drugs calculated accordingly. Dose response curves were plotted in Graph-pad Prism.

### PF culture and the SYBR Green I^®^ assay for parasite growth inhibition

PF 3D7 or C580Y lines were cultured and maintained at 1-5% parasitaemia in fresh group O-positive red blood cells re-suspended to a 5% haematocrit in custom reconstituted RPMI 1640 complete media (Thermo Scientific) containing 0.23% sodium bicarbonate, 0.4% D-glucose, 0.005% hypoxanthine 0.6% Hepes, 0.5% Albumax II, 0.03% L-glutamine and 25mg/L gentamicin. Culture flasks were gassed with a mixture of 1% O2, 5% CO2, and 94% N2 and incubated at 37°C. Prior to the start of the experiments, asynchronous stock cultures containing mainly ring stages were synchronised with 5% sorbitol as previously described. ^59^ Parasitaemia was determined with drug assays performed when the parasitaemia was between 1.5-5% with >90% rings. The stock culture was diluted to a haematocrit of 4% and 0.3% parasitaemia in complete media following which 50µl was mixed with 50µl of serial diluted drugs/inhibitors in complete media pre-dispensed in black 96 well optical culture plates (Thermo scientific) for a final haematocrit of 2%. Plates were gassed and incubated at 37° C for 72 hours followed by freezing at −20° C for at least 24 hours. The plate setup also included no drug controls as well as uninfected red cells at 2% haematocrit. After 72 hours of incubation and at least overnight freezing at −20° C, plates were thawed at room temperature for ∼4 hours. This was followed by addition of 100µl to each well of 1X SYBR Green I® (Invitrogen) lysis buffer containing 20 mM Tris, 5 mM EDTA, 0.008% saponin and 0.08% Triton X-100. Plate contents were mixed thoroughly by shaking at 700 rpm for 5 minutes and incubated for 1 hour at room temperature in the dark. After incubation, plates were read to quantify SYBR Green I® fluorescence intensity in each well by a PHERAstar® FSX microplate reader (BMG Labtech) with excitation and emission wavelengths of 485 and 520nm respectively. To determine growth inhibition, background fluorescence intensity from uninfected red cells was subtracted first. Fluorescence intensity of no drug controls was then set to correspond to 100% and subsequent intensity in presence of drug/inhibitor was calculated accordingly. Dose response curves and IC_50_ concentrations were plotted in Graph-pad Prism 7. Human blood was obtained and used within the ethical remit of the Scottish National Blood Transfusion Service.

### *In vitro* drug combinations

Parasites were maintained and cultivated as described above. To determine drug interactions of DHA in combination with DUB or proteasome inhibitors, serial dilutions of DHA were mixed with fixed ratios of epoxomicin, b-AP15, PR-619 and WP1130 or their fractional combinations at their respective IC_50_s or half IC_50_s. The drug combinations were incubated with parasites from which parasite growth was quantified and dose response curves were plotted, for DHA alone or in combination with the fixed doses of the DUB or proteasome inhibitors. IC_50_ values were obtained and the fold change or IC_50_ shifts were plotted in Graph-pad Prism using the extra sum of squares F-test for statistical comparison. For drug interactions in fixed ratios, a modified fixed ratio interaction assay was employed as previously described. ^60^ Drug combinations were prepared in six distinct molar concentration combination ratios; 5:0, 4:1, 3:2, 2:3 1:4, 0:5 and dispensed in top wells of 96-well plates. This was followed by a 2 or 3-fold serial dilution with precisely pre-calculated estimates that made sure that the IC_50_ of individual drugs falls to the middle of the plate. The drug combinations were then incubated with parasites from which parasite growth and dose response curves were calculated for each drug alone or in combination. Fractional inhibitory concentrations (FIC_50_) were obtained for drugs in combination and summed to obtain the ∑FIC_50_ using the formula below:

∑FIC_50_ = (IC_50_ of drug A in combination/ IC_50_ of drug A alone) + (IC_50_ of drug B in combination/ IC_50_ of drug B alone).

An ∑FIC_50_ of >4 was used to denote antagonism, ∑FIC_50_ ≤0.5 synergism and ∑FIC_50_ = 0.5-4 additivity. ^61^ FIC_50_ for the drug combinations were plotted to obtain isobolograms for the drug combination ratios.

### PF viability assays

The 3D7 line was synchronised with 5% sorbitol over three life cycles followed by Nycodenz enrichment of later schizonts. Enriched schizonts were incubated with fresh red blood cells in a shaking incubator for 3 hours followed by another round of sorbitol treatment to eliminate residual late stage parasites. Resultant ring cultures were diluted to around ∼1% parasitaemia and incubated with predefined drug combinations for set time periods. Drugs were washed off 3 times after the set incubation times. Parasite viability was assessed 66 hours later in cycle 2 by flow cytometry analysis of parasite cultures stained with Syber Green I and MitoTracker Deep Red dyes (Invitrogen). Flow cytometry analysis was carried on a MACSQuant® Analyzer 10.

### *In vivo* anti-parasitic activity of DUB inhibitors

To evaluate the activity of DUB inhibitors (b-AP15) *in vivo*, the Peters’ 4 day suppressive test was initially employed as previously described. ^62^ Stock concentrations of b-AP15 were prepared at 3mg/ml and 1mg/ml in a 1:1 mixture of DMSO and Tween^®^ 80 (Sigma) followed by a 10-fold dilution to stock working concentrations (5% DMSO and Tween^®^ 80 final) in sterile distilled water. CQ was prepared at 50mg/ml in 1X PBS and diluted to working stock in 1X PBS. A donor mouse was initially infected with PB 820 line from which blood was obtained when the parasitaemia was between 2-5%. Donor blood was diluted in rich PBS following which ∼10^5^ parasites were inoculated by IP into four mice groups (3 mice per group). 1-hour post infection, mice groups received drug doses by IP injection as follows: group 1 (vehicle; 5% DMSO & Tween^®^ 80), group 2 (CQ; 20mg/kg), group 3 (b-AP15; 1mg/kg) and group 4 (5mg/kg) for 4 consecutive days. Parasitaemia was monitored daily by flow cytometry analysis of infected cells stained with Hoechst-33258 and microscopic analysis of methanol fixed Giemsa stained smears. To evaluate the potential synergy of b-AP15 and ART *in vivo*, a modified Rane’s curative test in established infections was used. ^63^ Blood was obtained from a donor mouse at a parasitaemia of 2-3% and diluted in rich PBS. Seventeen mice were inoculated with ∼10^5^ parasites by IP on day 0 allowing the parasitaemia to rise to ∼2-2.5%, typically on day 4. Following the establishment of infection, mice were divided into five groups and received drug doses as follows: group 1 (5mg/kg ART n=3), group 2 (10mg/kg ART, n=3), group 3 (20mg/kg ART n=3), group 4 (5mg/kg ART + 1mg/kg b-AP15, n=4), group 5 (10mg/kg ART + 1mg/kg b-AP15, n=4). ART and b-AP15 were prepared at 12.5mg/ml and 1mg/ml respectively in 1:1 mixture of DMSO and Tween^®^ 80 and diluted 10-fold (final 5% DMSO and Tween^®^ 80) to their respective working concentrations. Parasitaemia was monitored daily by flow cytometry and analysis of methanol fixed Giemsa stained smears.

## Acknowledgments

We thank Mathias Matti for meaningful discussions; Steve Ward at Liverpool School of Tropical Medicine for suggestions and carefully reading the manuscript; Diane Vaughan and the iii-flow cytometry facility for technical assistance. We also thank Amit Mahindra and Andrew Jamieson at the University of Glasgow School of Chemistry for HPLC analysis of the DUB inhibitors. This work was supported by grants from the Wellcome Trust to A.P.W (083811/Z/07/Z; 107046/Z/15/Z). M.P.B. and APW are funded by a Wellcome Trust core grant to the Wellcome Centre for Integrative Parasitology (104111/Z/14/Z). N.V.S is a Commonwealth Doctoral Scholar (MWCS-2017-789), funded by the UK government.

## Author contributions

N.V.S conceived the experiments, performed data curation, analysis, investigation, validation, visualisation and writing of original draft. K.R.H, M.T.R and M.P.B participated in formal data analysis, investigation, validation, review and editing. A.P.W conceived the study, experiments, analysis, investigation, validation, writing of original draft, review, editing and supervision.

## Notes

### Competing Interest Statement

The authors have declared no competing interest.

