## Supplementary materials for "Mammalian deubiquitinating enzyme inhibitors display *in vitro* and *in vivo* activity against malaria parasites and potentiate artemisinin action"

**Supplementary table and figure legends**

**Supplementary Table 1: A manually created list of 17 DUBs in malaria parasites, their predicted function and essentiality.** The list was created based on previous *in silico* predictions [1-2] and a selected functional studies [3-4] as well as recent genome wide knockout screens. [5-6]

**Supplementary Table 2: A list of DUB inhibitors used in the study, their targets, chemical structure, supplier and primary references.**

**Supplementary Table 3: HPLC chemical purity and retention times of DUB inhibitors used in this study**

**Supplementary Figure 1: UCH-L3 inhibitor displays no activity in malaria parasites. A, B.** A selected growth inhibition plots of TCID in 3D7 (**A**) and 820 line (**B**). **C.** Amino acid sequence alignment of indicated *Plasmodium* spp. UCH-L3 against human UCH-L3. Conserved residues across human and *Plasmodium* UCH-L3s are indicated by asterisks. **D**. Phylogenetic tree of human, mouse and *Plasmodium* UCH-L3 predicted protein sequences showing their evolutionary divergence**.**

**Supplementary Figure 2: b-AP15 potentiates DHA action in UBP-1 G1808$^{V2721F}$ and G1807$^{V2752F}$ mutant lines**. **A, B**. Dose response curves and $IC_{50}$ value of DHA alone or combined with b-AP15 at $IC_{50}$ (DHA δ) in the UBP-1 G1808$^{V2721F}$ (**A**) and G1807$^{V2752F}$ UBP-1 mutant lines.

**Supplementary Figure 3:**

**A, B**. Dose response curves and $IC_{50}$ values of DHA alone or combined with WP1130 (DHA ε) (**A**) or PR-619 (DHA λ) (**B**) at $IC_{50}$ concentration in the PF 3D7 line. **C, D**. Isobologram plots of DHA in combination with b-AP15 **(C)** and WP1130 **(D)** and their raw $\sum FIC_{50}$ values. $\sum FIC_{50}$ values, plotted $FIC_{50}$s and error bars are means and standard deviations from three biological repeats.

**Supplementary Figure 4:**

Amino acid sequence alignment of indicated PB and PF USP14. Conserved residues are indicated by asterisks.

**Supplementary Figure 5**

HPLC chromatograms of the drug solvent (DMSO) and indicated DUB inhibitors. HPLC runs were carried out as described in methods.

**List of Supplementary Figures**

**Supplementary Table 1**

| PF/PB gene ID | Close human orthologue | Essential? (PF/PB) | Predicted function in malaria parasites |
|---|---|---|---|
| PF3D7_1460400/PBANKA_1324100 | UCH-L3 | Yes/not characterised | deNeddylase/DUB activity |
| PF3D7_1117100/PBANKA_0930900 | UCH54 | Yes/dispensable | deNeddylase/DUB activity |
| PF3D7_0726500/PBANKA_0210600 | UCH-L1 | Yes/Yes | Not known |
| PF3D7_0104300/PBANKA_0208800 | HAUSP/USP7 (UBP-1) | Yes/not characterised | Implicated in drug resistance |
| PF3D7_0413900/PBANKA_0715900 | USP13 | Yes/not characterised | Not known |
| PF3D7_0527200/PBANKA_1242000 | USP14 | Yes/dispensable | DUB activity |
| PF3D7_0516700/PBANKA_1231500 | USP2 | not characterised/dispensable | Not known |
| PF3D7_0904600/PBANKA_0416800 | USP14? | dispensable/dispensable | Not known |
| PF3D7_1317000/PBANKA_1415500 | USP39 | Yes/not characterised | Not known |
| PF3D7_1414700/PBANKA_1028000 | UCH36? | dispensable/not characterised | Not known |
| PF3D7_1226800/PBANKA_1441600 | Ataxin 3 | Yes/dispensable | Not known |
| PF3D7_0403500/PBANKA_1001100 | USP? | Yes/Yes | Not known |
| PF3D7_1111900/PBANKA_0935700 | Josephin domain | Yes/not characterised | Not known |
| PF3D7_0923100/PBANKA_0824000 | OTU domain | dispensable/dispensable | Not known |
| PF3D7_1031400/PBANKA_0515350 | OTU like | not characterised/dispensable | Not known |
| PF3D7_1141700/PBANKA_0907300 | OTU domain | Yes/not characterised | Not known |
| PF3D7_0920300/PBANKA_0821200 | OTU domain | Yes/not characterised | Not known |

**Supplementary Table 2**

| Inhibitor, Mw | UPS target | Supplier | Purity grade | Chemical structure |
|---|---|---|---|---|
| PR-619, 223.28 | broad spectrum DUB inhibitor[a] | Focus biomolecules (CAS #: 2645-32-1) | 98% by TLC NMR | 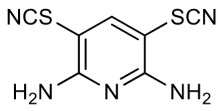 |
| P5091, 348.23 | USP7 and USP47 DUBs[b] | Focus biomolecules (CAS #: 882257-11-6) | 98% by TLC NMR | 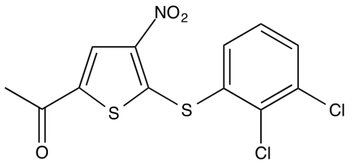 |
| TCID, 283.93 | UCH-L3 and UCH-L1 DUBs[c] | Focus biomolecules (CAS #: 30675-13-9) | 97% by TLC NMR | 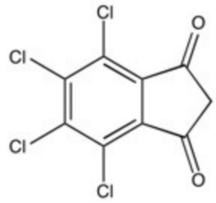 |
| WP1130 | UCH-L1, USP9X, USP14, UCH37 DUBs[d] | Focus biomolecules (CAS #: 856243-80-6) | 98% by TLC NMR | 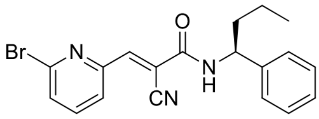 |
| b-AP15, 419.4 | USP14 and UCH-L5 DUBs[e] | Focus biomolecules (CAS #: 1009817-63-3) | >98% by HPLC | 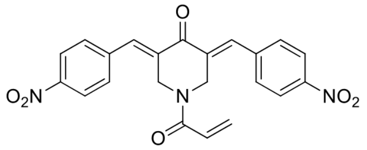 |
| NSC-632839, 339.86 | USP2, USP7, SENP2 DUBs[f] | Focus biomolecules (CAS #: 157654-67-6) | >98% by HPLC NMR | 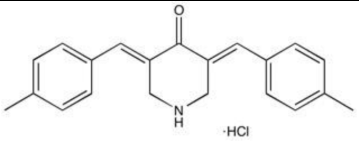 |
| 1,10 phenanthroline, 198.2 | Metalloproteases and JAMM isopeptidases[g] | BPS biosciences (CAS #: 5144-89-8) | ≥99% by HPLC | 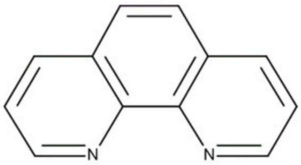 |

a, [7] b,[8] c, [9] d, [10] e, [11] f, [12] g. [13]

**Supplementary Table 3**

| DUB inhibitor | HPLC Purity (%) | Retention time (minutes) |
|---|---|---|
| 1,10-phenanthroline | >99 | 12.433 |
| b-AP15 | >99 | 19.095 |
| P5091 | >99 | 21.303 |
| PR-619 | >99 | 13.708 |
| WP1130 | >99 | 20.205 |
| TCID | >99 | 19.755 |
| NSC-632839 | 99 | 16.612 |

**Supplementary Figure 1**

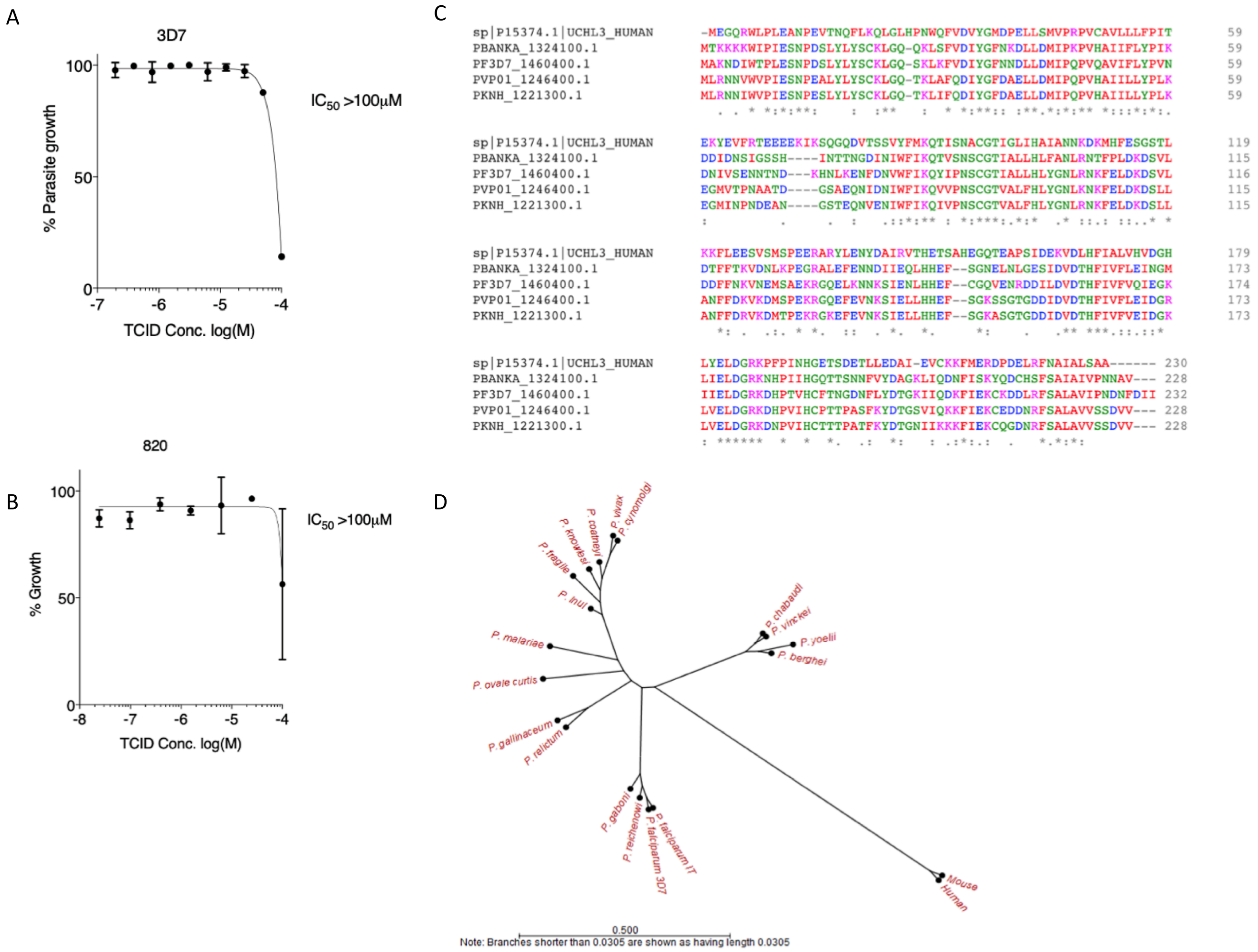

**Supplementary Figure 2**

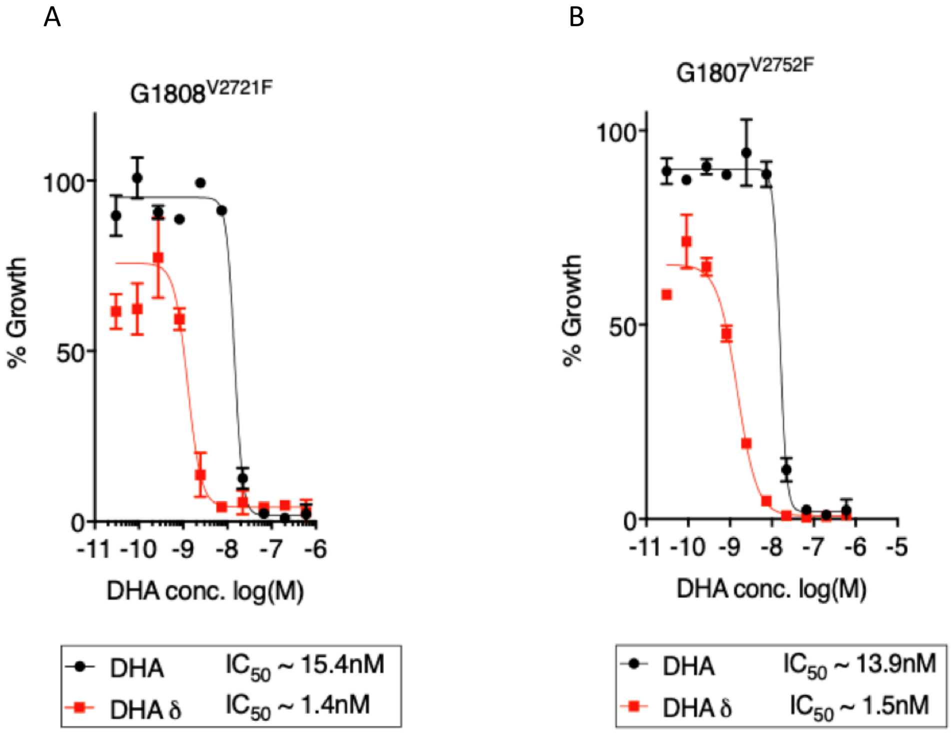

**Supplementary Figure 3**

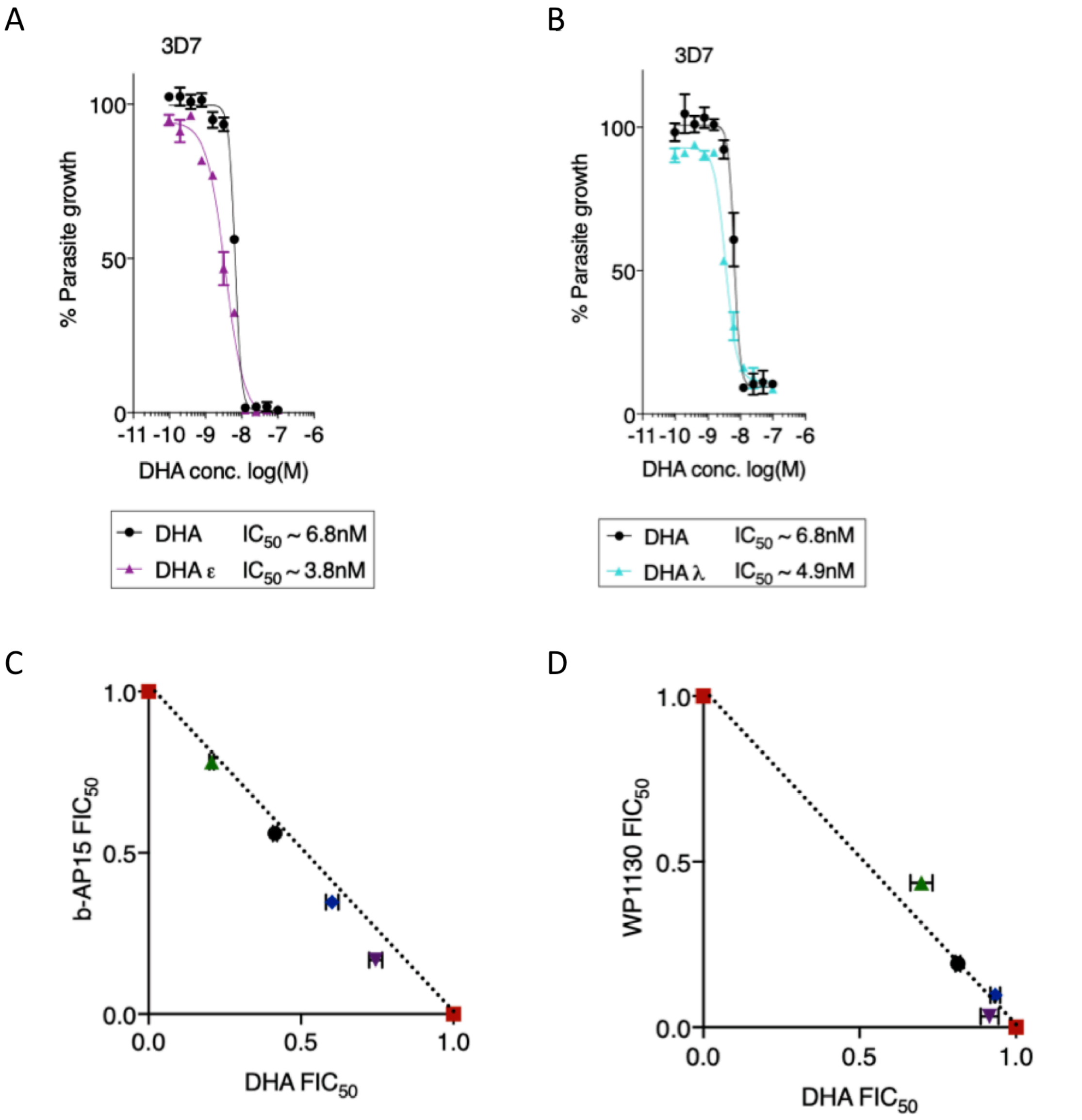


| Drug Combination Ratio | $\sum FIC_{50}$ |
| --- | --- |
| DHA 100% | 1 |
| DHA 80% : b-AP15 20% | 0.891 |
| DHA 60% : b-AP15 40% | 0.973 |
| DHA 40% : b-AP15 60% | 0.983 |
| DHA 20% : b-AP15 80% | 1.02 |
| b-AP15 100% | 1 |

| Drug Combination Ratio | $\sum FIC_{50}$ |
| --- | --- |
| DHA 100% | 1 |
| DHA 80% : WP1130 20% | 0.924 |
| DHA 60% : WP1130 40% | 1.032 |
| DHA 40% : WP1130 60% | 1.004 |
| DHA 20% : WP1130 80% | 1.094 |
| WP1130 100% | 1 |

**Supplementary Figure 4**

```
PF3D7_0527200.1      MTLVNITVKWKNQVFNNIELDVSEPLILLKTQLWQLTNVPPEKQKLMYKGLLKDDVDLSL     60
PBANKA_1242000.1     MTIVKVTVKWKTNIYNDLELNINEPIALFKEQLWKITSVPPEKQKLMYKGLIKNDTNLHT     60
                     **:*::*****.:::*::**::.**: *:* ***::*.*************:*:*.:*

PF3D7_0527200.1      LNIKENDKIMLVGSAESLVEKPKDIIFEEDLTNEEKQKIHTKENIIFEEQGIVNLGNTCY     120
PBANKA_1242000.1     LNIKNNDKIMLVGSSELLAEKPDKITFFEDLSKEDKEKLNEDKNIIFEEQGIVNLGNTCY     120
                     ****:*********:* *.***..* * ***::*:*:*:: .:****************

PF3D7_0527200.1      FNAVLQFLTSFDDLGNFLRSYKSKESKLIKTNKDILFDSFIEFAHSFEKSSEPYVPVTLL     180
PBANKA_1242000.1     FNAVLQFLTSFNDLGEFLSNIKKTQNFGFRSNKDILFDCYIHFSQTFGKSSKPYVPLELL     180
                     ***********:***:** . *..:.  :::*******.:*.*:::* ***:****: **

PF3D7_0527200.1      KSFRDVYPKFKSVNLRTKQYAQQDAEECMNAILTCLNEQTDNKIIDKLFSFQIISNMKFV     240
PBANKA_1242000.1     KAFRDVFPKFKTINIRTKQYAQQDAEECMNAILTSLNDHTESKIIDKLFSFKIIGKIKCV     240
                     *:****:****::*:******************.**::*:.*********:**.::* *

PF3D7_0527200.1      ETVEQHEQKEKKDEKKDEKKDEKKDEKKEEKKEEIKNINITSNNNNIDNNNDNNDNNNNN     300
PBANKA_1242000.1     EQNSQDDEPNKQKESQK---NENSQ-----------------------------------     262
                     *  .*.:: :*:.*.:.   :*:.:

PF3D7_0527200.1      NNNSVQQNDHNNKDISHNNIFETTQEFNNKLICYMGTPNTPVNHLHEGIRLSLHEKIRKN     360
PBANKA_1242000.1     ---KPESDKNNPSQNEENDHFELTQEKHNKLICYMGTQNTPVNHLHEGIRLSLIEKIKKK     319
                        . :.:.:* .: ..*: ** *** :********* ************** ***:*:

PF3D7_0527200.1      RNED-NKECIYEKKSEINSLPPYLIVHFLRFESKKIVESNNSGVSVVTAKICRKVSFPDT     419
PBANKA_1242000.1     KNDNDKEDTLYEKKSEIDSLPPYLIIHFLRFESKRIVDTSN-TVSVVTAKICRKVSFPEI     378
                     :*:: ::: :*******:*******:********:**::.*  ***************:

PF3D7_0527200.1      FDMYDFCSEKIKEELKIARDIIMKRKDKETSLSPQKENIQNIEQNNINNQYNQNNSNDNP     479
PBANKA_1242000.1     FDIYDFCSDRIKADLKISRNIIMNRKDIKTPIVQENKNEDD-------------------     419
                     **:*****::** :***:*:***:*** :* :  :::* ::

PF3D7_0527200.1      KHVQEHT-NQINKEELIELPTGEYELISVITHKGRNEESGHYIAWKKMKKFFSSNSNIDQ     538
PBANKA_1242000.1     KMLETLTNQDDKNKEFVEIPNGEYELISVITHKGRNEESGHYIAWKKMRNRVNKNSEYDA     479
                     * ::  * :: :::*::*:*.***************************:: ...**: *

PF3D7_0527200.1      NESSNKKTKNANDSLWLKMDDDKVSTHKFSSIDFYGGCSDYNIAVLLLYKRKNISCTPDE     598
PBANKA_1242000.1     NGPRTKKNKSNNEPTWFKMDDDKVSLHNFSSLDLFGGCSDYNTAVLLLYKRKTISCTKDE     539
                     *   .**.*. *:  *:******** *:***:*::******* *********.**** **

PF3D7_0527200.1      MNMDIKE   605
PBANKA_1242000.1     LNKYSV-   545
                     :*
```

**Supplementary Figure 5**

**HPLC chromatogram of DMSO**

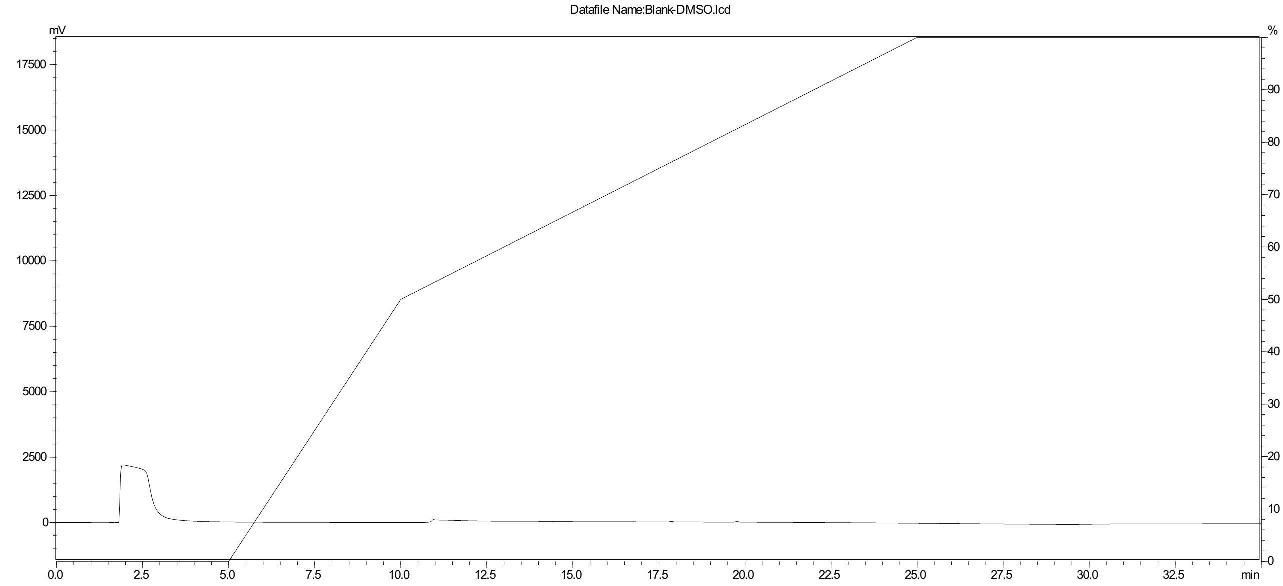


**HPLC chromatogram of 1,10-phenanthroline**

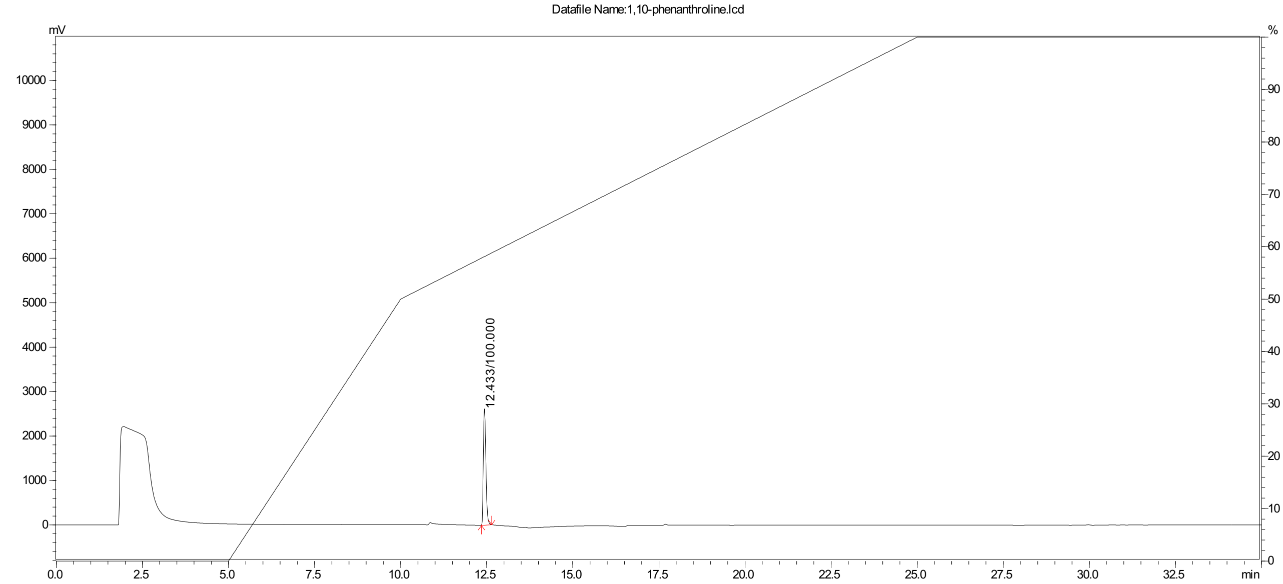

**HPLC chromatogram of 1,10-phenanthroline, gradient zoom**

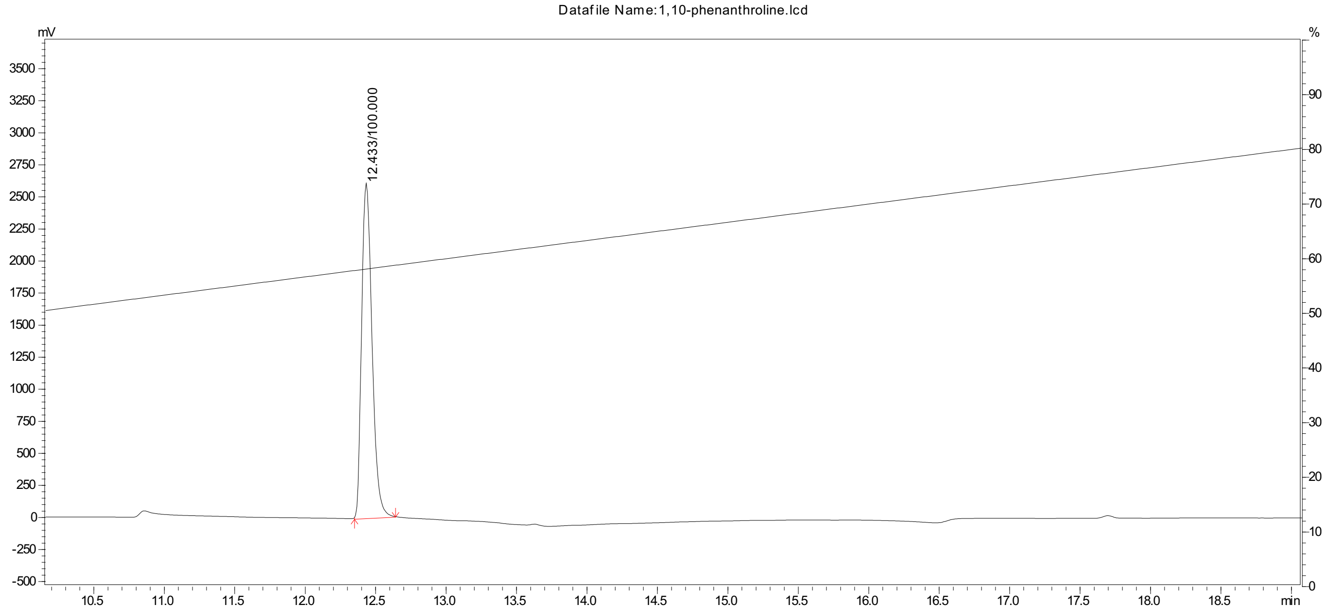


**HPLC chromatogram of b-AP15**

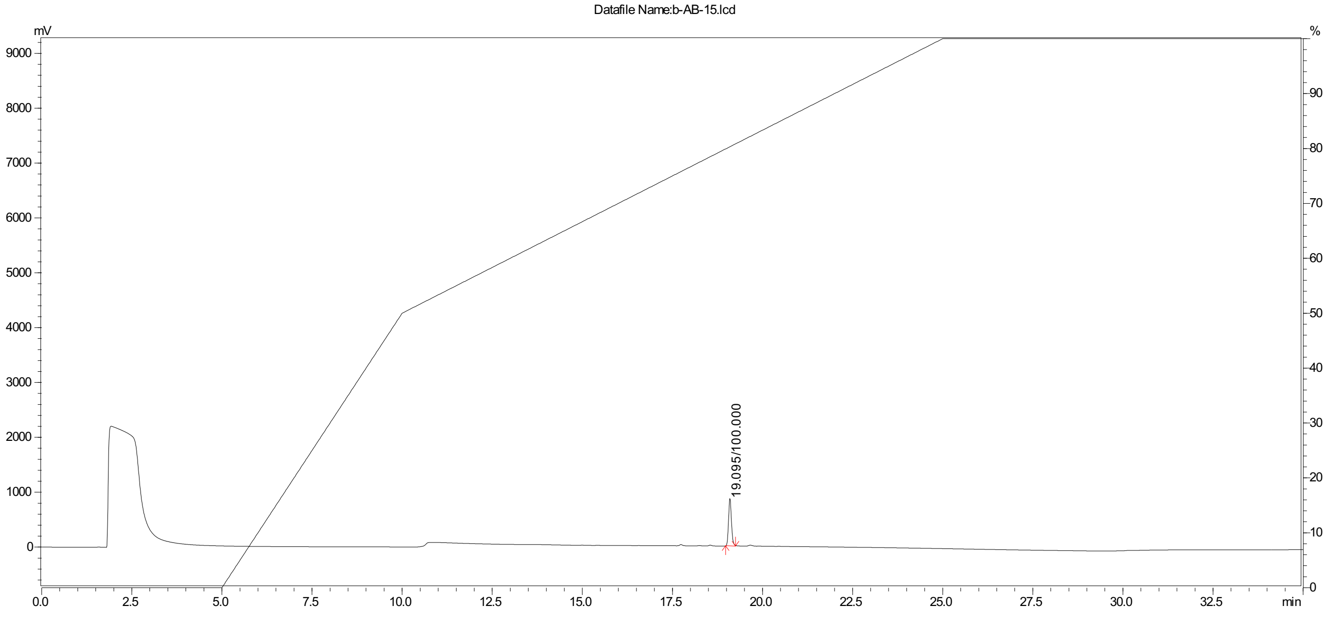

**HPLC chromatogram of b-AP15, gradient zoom**

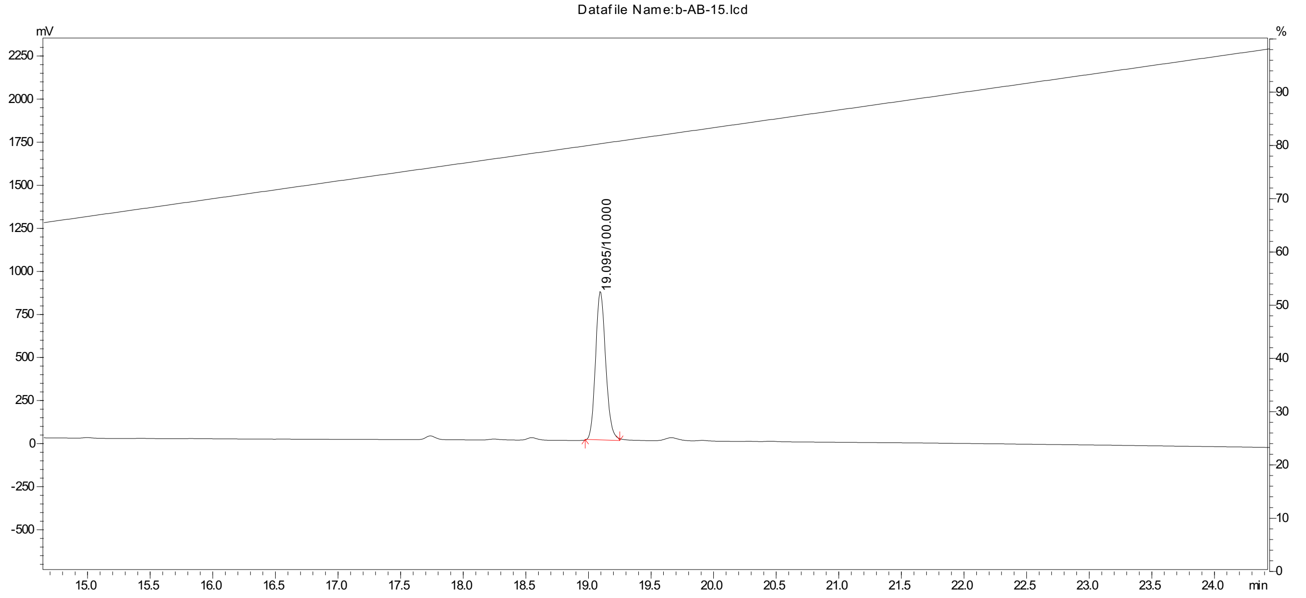


**HPLC chromatogram of P5091**

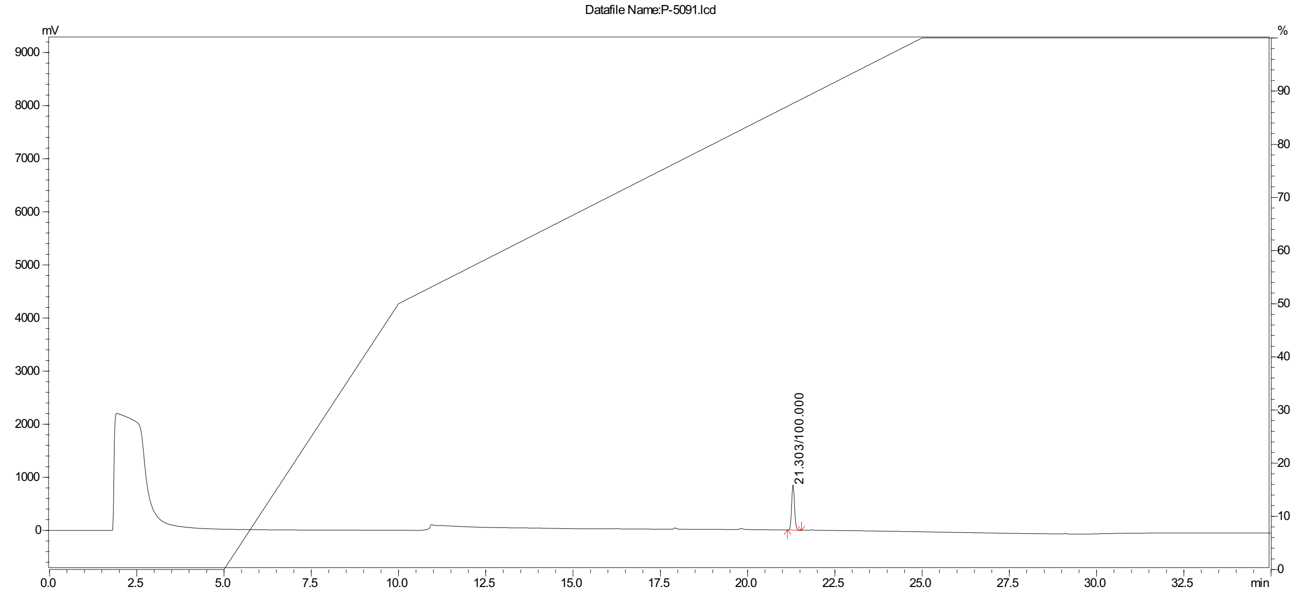

**HPLC chromatogram of P5091, gradient zoom**

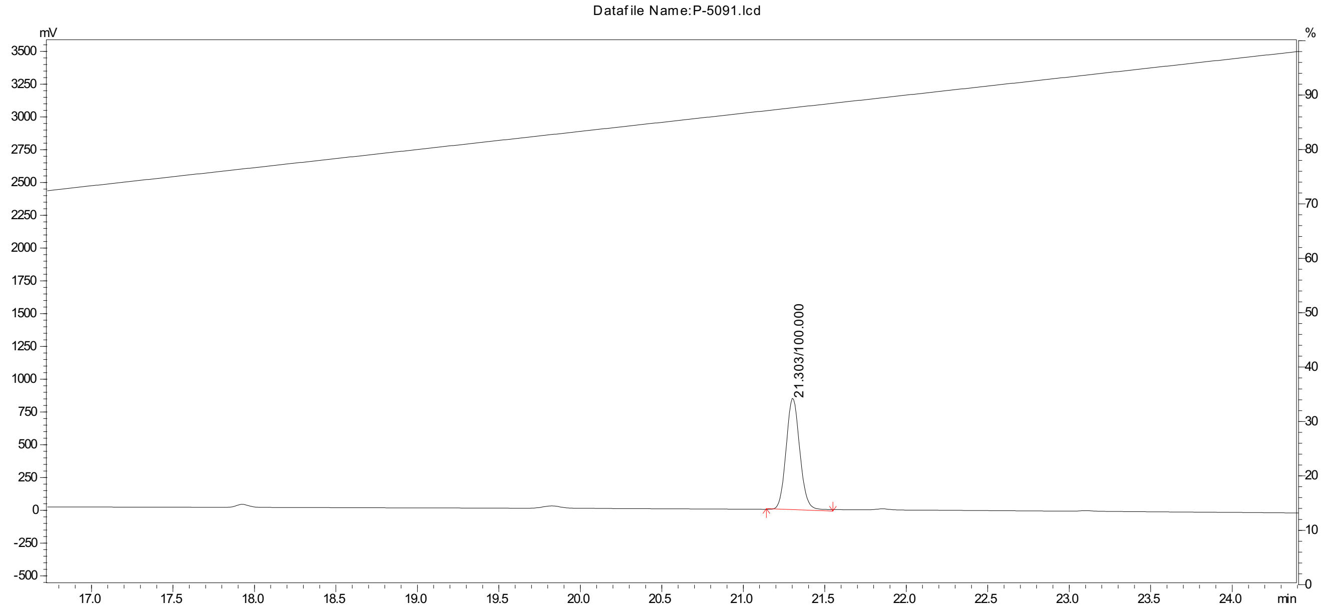


**HPLC chromatogram of PR-619**

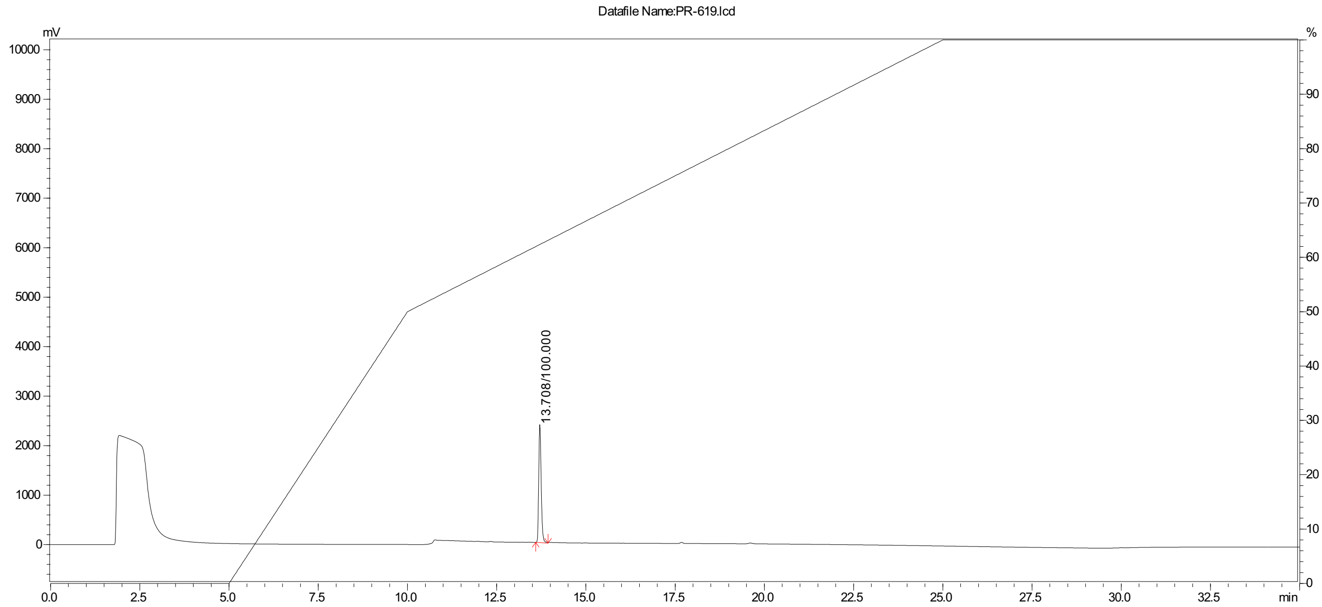

**HPLC chromatogram of PR-619, gradient zoom**

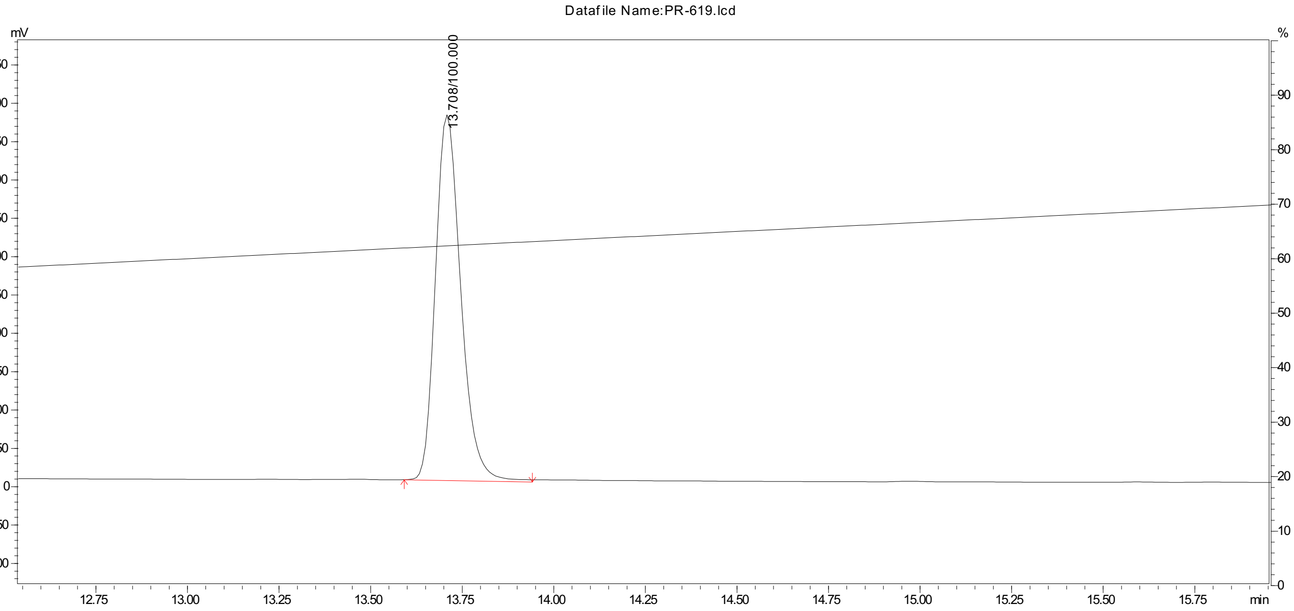


**HPLC chromatogram of WP-1130**

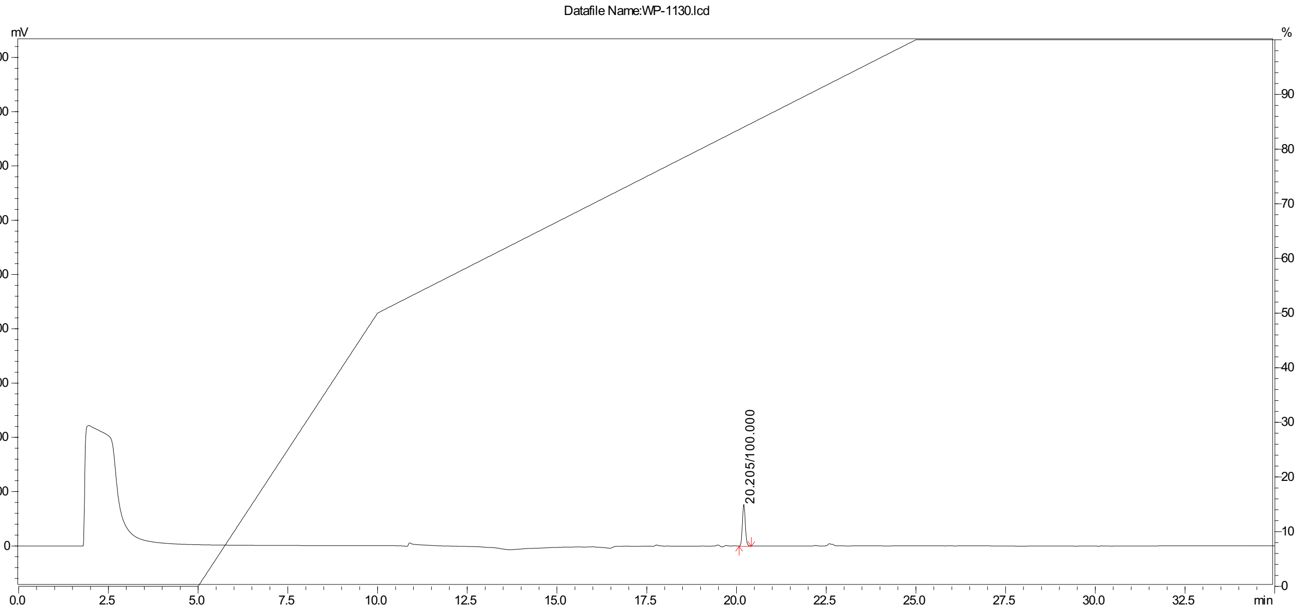

**HPLC chromatogram of WP-1130, gradient zoom**

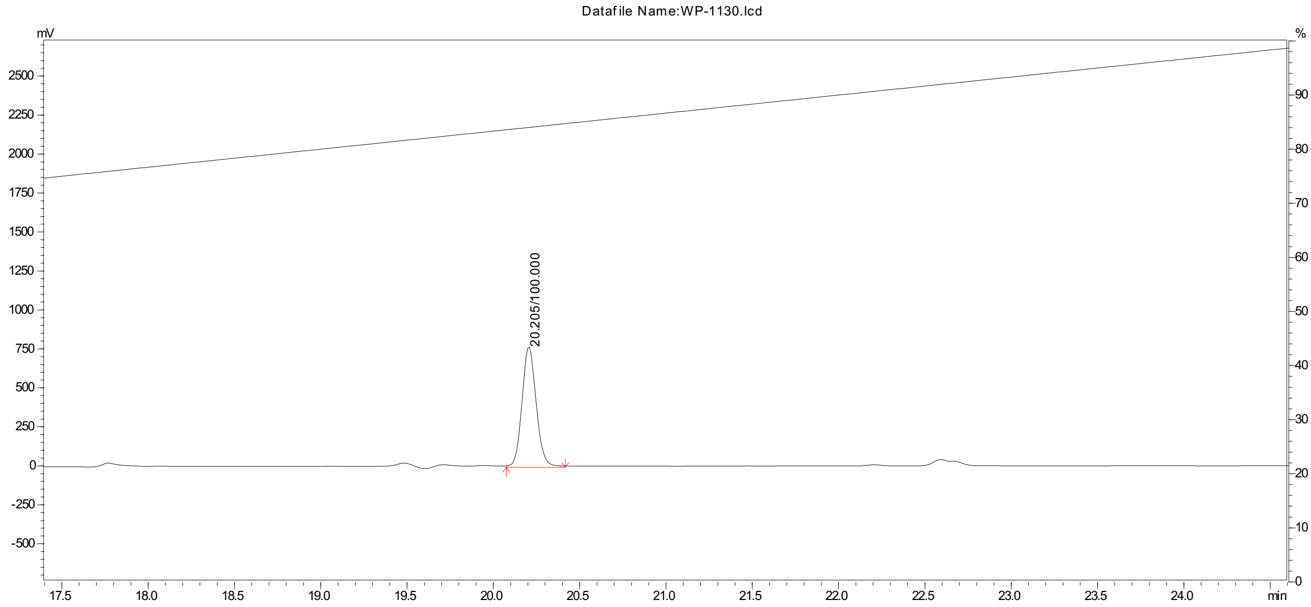


**HPLC chromatogram of TCID**

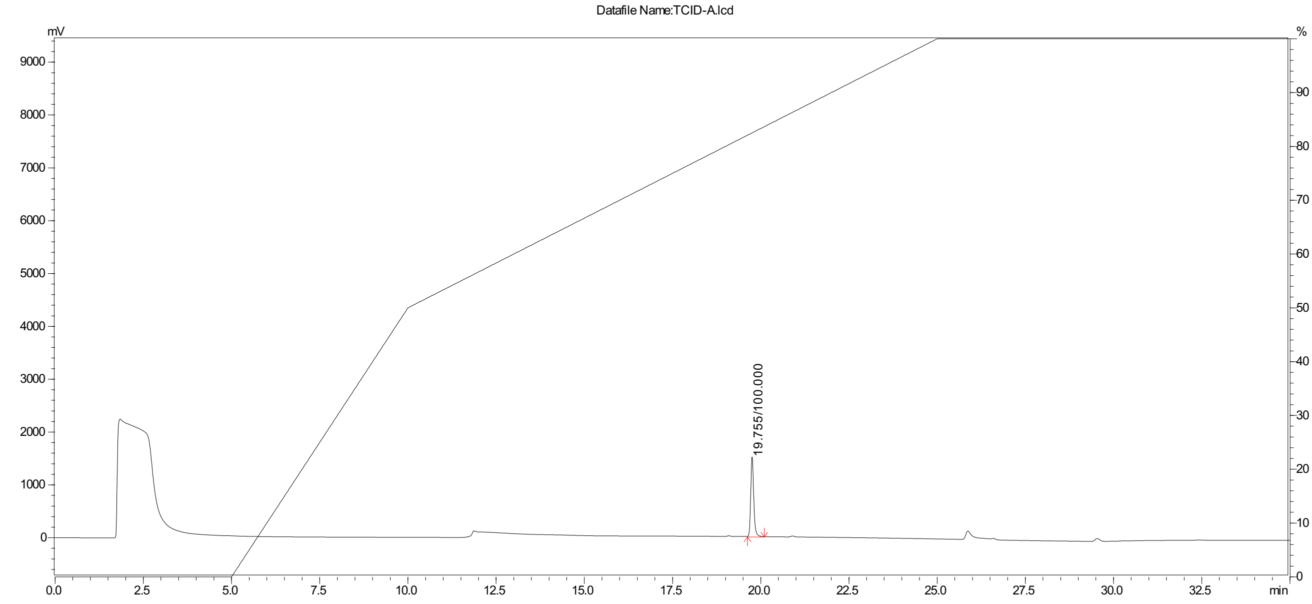

**HPLC chromatogram of TCID, gradient zoom**

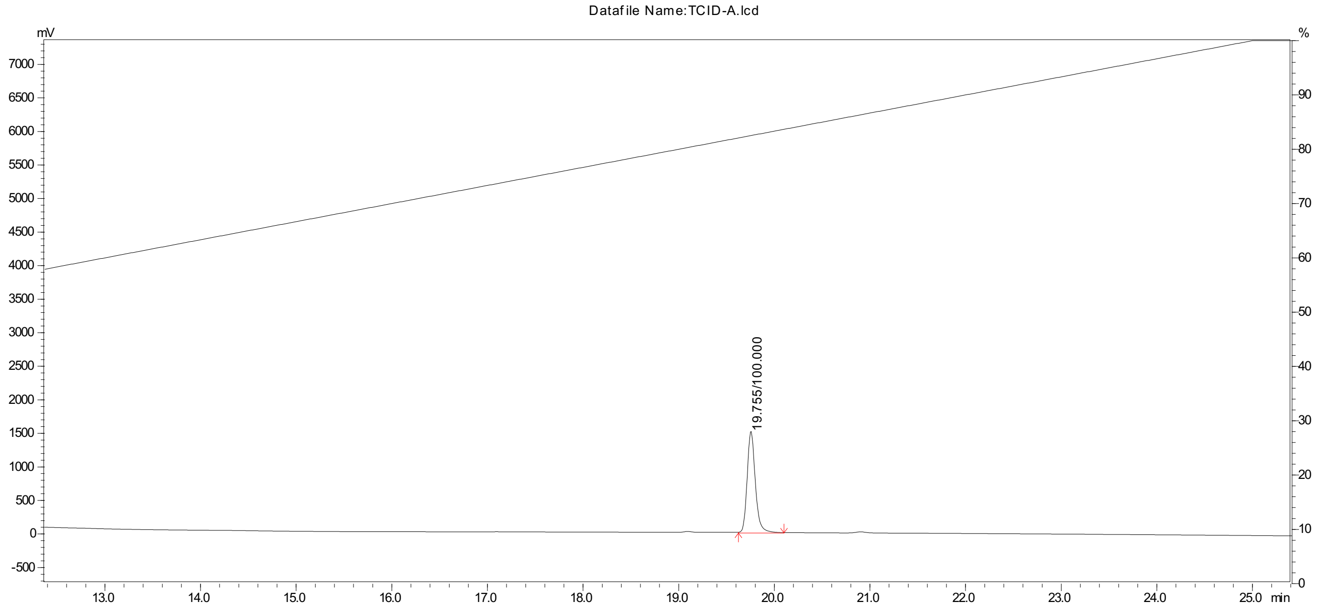


**HPLC chromatogram of NSC-632839**

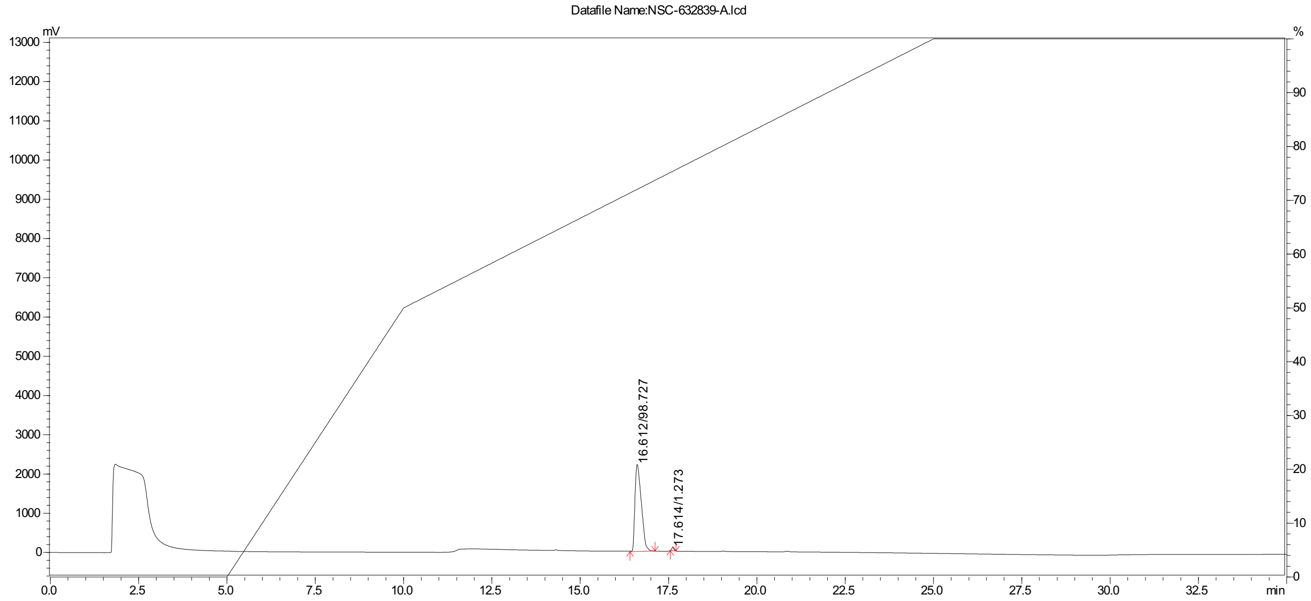

**HPLC chromatogram of NSC-632839, gradient zoom**

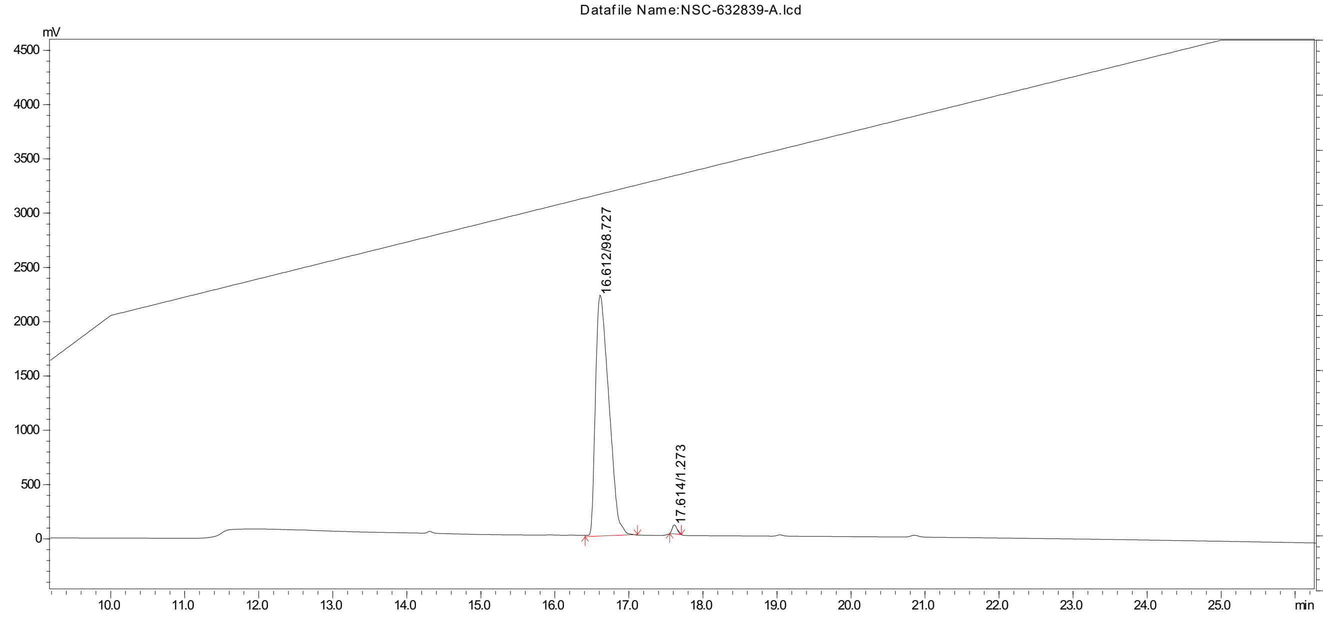
